# Identification of retinal tau oligomers, citrullinated tau, and other tau isoforms in early and advanced AD and relations to disease status

**DOI:** 10.1101/2024.02.13.579999

**Authors:** Haoshen Shi, Nazanin Mirzaei, Yosef Koronyo, Miyah R. Davis, Edward Robinson, Gila M. Braun, Ousman Jallow, Altan Rentsendorj, V Krishnan Ramanujan, Justyna Fert-Bober, Andrei A. Kramerov, Alexander V. Ljubimov, Lon S. Schneider, Warren G. Tourtellotte, Debra Hawes, Julie A. Schneider, Keith L. Black, Rakez Kayed, Maj-Linda B. Selenica, Daniel C. Lee, Dieu-Trang Fuchs, Maya Koronyo-Hamaoui

## Abstract

**Importance:** This study identifies and quantifies diverse pathological tau isoforms in the retina of both early and advanced-stage Alzheimer’s disease (AD) and determines their relationship with disease status.

**Objective:** A case-control study was conducted to investigate the accumulation of retinal neurofibrillary tangles (NFTs), paired helical filament (PHF)-tau, oligomeric tau (oligo-tau), hyperphosphorylated tau (p-tau), and citrullinated tau (Cit-tau) in relation to the respective brain pathology and cognitive dysfunction in mild cognitively impaired (MCI) and AD dementia patients versus normal cognition (NC) controls.

**Design, setting and participants:** Eyes and brains from donors diagnosed with AD, MCI (due to AD), and NC were collected (n=75 in total), along with clinical and neuropathological data. Brain and retinal cross-sections–in predefined superior-temporal and inferior-temporal (ST/IT) subregions–were subjected to histopathology analysis or Nanostring GeoMx digital spatial profiling.

**Main outcomes and measure:** Retinal burden of NFTs (pretangles and mature tangles), PHF-tau, p-tau, oligo-tau, and Cit-tau was assessed in MCI and AD versus NC retinas. Pairwise correlations revealed associations between retinal and brain parameters and cognitive status.

**Results:** Increased retinal NFTs (1.8-fold, p=0.0494), PHF-tau (2.3-fold, p<0.0001), oligo-tau (9.1-fold, p<0.0001), CitR_209_-tau (4.3-fold, p<0.0001), pSer202/Thr205-tau (AT8; 4.1-fold, p<0.0001), and pSer396-tau (2.8-fold, p=0.0015) were detected in AD patients. Retinas from MCI patients showed significant increases in NFTs (2.0-fold, p=0.0444), CitR_209_-tau (3.5-fold, p=0.0201), pSer396-tau (2.6-fold, p=0.0409), and, moreover, oligo-tau (5.8-fold, p=0.0045). Nanostring GeoMx quantification demonstrated upregulated retinal p-tau levels in MCI patients at phosphorylation sites of Ser214 (2.3-fold, p=0.0060), Ser396 (1.8-fold, p=0.0052), Ser404 (2.4-fold, p=0.0018), and Thr231 (3.3-fold, p=0.0028). Strong correlations were found between retinal tau forms to paired-brain pathology and cognitive status: a) retinal oligo-tau vs. Braak stage (r=0.60, P=0.0002), b) retinal PHF-tau vs. ABC average score (r=0.64, P=0.0043), c) retinal pSer396-tau vs. brain NFTs (r=0.68, P<0.0001), and d) retinal pSer202/Thr205-tau vs. MMSE scores (r= –0.77, P=0.0089).

**Conclusions and Relevance:** This study reveals increases in immature and mature retinal tau isoforms in MCI and AD patients, highlighting their relationship with brain pathology and cognition. The data provide strong incentive to further explore retinal tauopathy markers that may be useful for early detection and monitoring of AD staging through noninvasive retinal imaging.

## 1. Introduction

Alzheimer’s disease (AD) is the leading cause of senile dementia worldwide. The pathological hallmarks of AD are characterized by the accumulation of amyloid beta-protein (Aβ) deposits and abnormal microtubulin-associated tau protein aggregates in the brain ^1^. During the pathogenesis of AD, the tau protein undergoes post-translational modifications, such as hyperphosphorylation (p-tau) or citrullination (Cit-tau), which have an increased tendency to create toxic oligomers (oligo-tau) that propagate to healthy neurons and spread the disease ^2-4^. As the disease progresses, pathological tau species also aggregate into fibrils and paired helical filaments (PHF), ultimately forming intraneuronal neurofibrillary tangles (NFTs) ^5^. Both PHF-tau and NFTs can damage cytoplasmic functions and interfere with axonal transport, leading to neuronal cell death ^6^. While cerebral Aβ accumulation occurs decades prior to the manifestation of clinical symptoms ^7-9^, the subsequent increase in abnormal tau strongly associates with neurodegeneration and disease progression ^9-11^. The extended preclinical phase of AD allows for earlier, more effective intervention before severe neuronal loss occurs. Indeed, recent studies indicate enhanced efficacy for amyloid immunotherapies in early AD patients ^12,13^ and in individuals with a low brain tau burden, emphasizing the urgency to develop feasible, affordable, and non-invasive techniques for early screening and monitoring of AD.

The retina, an extension of the brain and the only central nervous system (CNS) organ not shielded by bone ^14,15^, holds promise for revolutionizing AD screening. Current limitations in early diagnosis and monitoring of AD in clinical settings ^16,17^ make retinal imaging a potential solution due to its accessibility for noninvasive, repeated, low cost imaging at ultra-high spatial resolution. Our group and others have demonstrated the manifestation of AD pathological features in the retina of mild cognitively impaired (MCI) and AD patients, including Aβ plaques, vascular Aβ deposits, various p-tau forms, inflammation, vascular damage, and neurodegeneration ^18-30^. Examination of postmortem retinal tissues from AD patients detected total tau, 3- and 4-repeat tau, p-tau, and NFTs-like structures ^18-21,23,31,32^. Retinal p-tau Ser202/Thr205 and Thr217 burdens correlated with Braak stage and p-tau Ser202/Thr205 burden in AD brains ^21^. Additionally, the severity stages of retinal p-tau Ser202/Thr205 correlated with Aβ phases in AD brains ^32^. Nevertheless, it remains unclear whether early and mature pathological tau forms exist and spread in the retina of AD patients at the earliest functional impairment (MCI) and dementia stages and, moreover, correlate with brain pathology and cognitive status.

To address these questions, we investigated aberrant tau forms associated with AD severity ^33^, including tau oligomers, p-tau, Cit-tau, PHF-tau, and NFTs in postmortem retinal tissues from a cohort of individuals with MCI (due to AD), AD dementia patients, and normal cognition (NC) controls. Our analyses indicate that most abnormal tau forms significantly increase in the retina of MCI and AD patients and generally correlate with the respective brain pathology. The results of this study may offer insights into the specific pathological tau forms that can serve as future retinal biomarkers for AD detection and assessment of disease progression.

## 2. METHODS

Extended methods are provided in Supplementary Materials.

### 2.1 Source of postmortem eyes and brains

Seventy-five eyes and 55 paired-brains from deceased AD dementia and MCI due-to-AD patients (n=45; neuropathologically confirmed as AD) and individuals with normal cognition (NC, n=30) were obtained from the University of Southern California Alzheimer’s Disease Research Center (USC-ADRC) Neuropathology Core (IRB-HS-042071), and the National Disease Research Interchange (NDRI, Philadelphia, PA) under Cedars-Sinai Medical Center IRB protocols, Pro00053412 and Pro00019393. Clinical and neuropathological data are detailed in **Table 1** and **Supplementary Table 1**. The available brain and retinal tissues for each tau marker analysis is specified in Table 1 for the histopathological cohort (NC: n=10-18 for each biomarker and 30 subjects in total, MCI: n=9-11 and AD: n=25-34) and for the NanoString GeoMx analysis cohort (NC: n=5-9, MCI: n=4-6, and AD: n=4-9).

**Table 1.** Demographic, neuropathological, and retinal tauopathy data on human donors in this study.

|  | NC |  |  | MCI |  |  | AD |  |  | Statistics |  |
| --- | --- | --- | --- | --- | --- | --- | --- | --- | --- | --- | --- |
|  | n | Mean | SD | n | Mean | SD | n | Mean | SD | F | P |
| Age (years) | 30 | 81.48 | 9.991 | 11 | 89.55 | 6.121 | 34 | 83.88 | 12.68 | 2.175 | 0.1212 |
| Gender | 17F (57%), 13M |  |  | 7F (64%), 4M |  |  | 21F (62%), 13M |  |  | - | - |
| Race (no.) | 25W (83%), 3H, 2B |  |  | 9W (82%), 1H, 1B |  |  | 24W (71%), 4A, 4H, 2B |  |  | - | - |
| PMI (h) | 27 | 7.44 | 3.526 | 11 | 8.182 | 4.899 | 34 | 8.529 | 4.203 | 0.543 | 0.5834 |
| MMSE Score | 4 | 26.25 | 2.754 | 5 | 21.8 | 4.207 | 10 | 12.7 | 8.982 | 6.103 | **0.0107** |
| CDR Score | 6 | 0.1667 | 0.4082 | 9 | 1.667 | 1.392 | 24 | 2.438 | 0.825 | 14.32 | **<0.0001** |
| Brain neuropathology |  |  |  |  |  |  |  |  |  |  |  |
| Braak Stage (%) | 10 | I-II (60%) |  | 11 | I-II (36%) |  | 34 | I-II (0%) |  | 21.69 | **<0.0001** |
|  |  | III-IV (30%) |  |  | III-IV (27%) |  |  | III-IV (24%) |  |  |  |
|  |  | V-VI (10%) |  |  | V-VI (36%) |  |  | V-VI (76%) |  |  |  |
| ABC (amyloid, Braak, CERAD) | 7 | 1.952 | 0.6506 | 9 | 2.148 | 0.58 | 31 | 2.733 | 0.3149 | 13.14 | **<0.0001** |
| Aβ plaque | 10 | 1.172 | 1.189 | 11 | 1.911 | 0.8236 | 34 | 2.53 | 0.9157 | 8.327 | **0.0007** |
| NFTs | 10 | 0.4965 | 0.5495 | 11 | 1.502 | 0.8077 | 34 | 2.363 | 0.8902 | 21.02 | **<0.0001** |
| NTs | 10 | 0.443 | 0.8439 | 11 | 1.001 | 0.824 | 34 | 1.78 | 1.178 | 6.966 | **0.0021** |
| Atrophy | 10 | 0.88 | 0.9897 | 11 | 1.114 | 0.9602 | 34 | 1.806 | 1.108 | 3.848 | **0.0276** |
| Retinal tau markers |  |  |  |  |  |  |  |  |  |  |  |
| pS202/T205-tau (AT8) | 18 | 1.35 | 1.439 | 8 | 4.683 | 4.78 | 20 | 4.08 | 3.796 | 3.348 | **0.0445** |
| pS212/T214-tau (AT100) | 9 | 13.69 | 6.507 | 4 | 18.08 | 9.629 | 6 | 21.04 | 5.426 | 2.104 | 0.1545 |
| pS396-tau (pS396) | 14 | 0.8023 | 0.4201 | 8 | 2.088 | 0.8238 | 25 | 2.223 | 1.433 | 7.441 | **0.0016** |
| CitR<sub>209</sub>-tau | 18 | 3.779 | 2.933 | 8 | 13.17 | 10.99 | 20 | 16.18 | 9.174 | 12.53 | **<0.0001** |
| Oligo-tau (T22) | 18 | 0.3124 | 0.2156 | 9 | 1.797 | 0.9237 | 19 | 2.833 | 1.523 | 25.7 | **<0.0001** |
| PHF-tau (PHF-1) | 9 | 392.3 | 155.1 | 5 | 383.5 | 188.5 | 10 | 941 | 234.4 | 20.35 | **<0.0001** |
| NFTs (MC-1) | 18 | 3.064 | 1.057 | 9 | 6.199 | 4.122 | 21 | 5.489 | 3.607 | 4.415 | **0.0178** |

List of human donors (total n = 75 subjects) included in this study. Paired brains with full neuropathological assessments were available for 55 human donors. Mean ABC scores were determined as follows: A, Aβ plaque score modified from Thal; B, NFT stage modified from Braak; C, neuritic plaque score modified from CERAD. Group values are presented as mean ± standard deviation (SD). *F* and *P*-values were determined using one-way analysis of variance with Tukey's multiple comparisons test. *P*-values presented in bold type demonstrate significance. Abbreviations: Aβ , amyloid beta-protein; AD, Alzheimer's disease; A, Asian; B, Black; CDR, Clinical Dementia Rating; CERAD, Consortium to Establish a Registry for Alzheimer's Disease; NC, normal cognition controls; F, female; H, Hispanic; M, male; MCI, mild cognitive impairment; MMSE, Mini-Mental State Examination; NFTs, neurofibrillary tangles; NTs, neuropil threads; PMI, post mortem interval; SD, standard deviation; W, White.

### 2.2 Clinical and neuropathological assessments

The USC-ADRC Clinical Core provided records of neurological examinations, cognitive evaluations, and extensive brain pathology reports. Neuropathological parameters were analyzed in multiple brain areas, including the hippocampus (particularly the Cornu ammonis CA1, at the level of the thalamic lateral geniculate body), entorhinal cortex, superior frontal gyrus of the frontal lobe, superior temporal gyrus of the temporal lobe, superior parietal lobule of the parietal lobe, primary visual cortex (Brodmann Area-17), and visual association (Area-18) of the occipital lobe. In all cases uniform brain sampling was done by a neuropathologist.

### 2.3 Eyes and brains processing and retinal and brain cross-section preparations for immunostaining, Gallyas and Hito Bielschowsky silver staining

Paired donor eyes and brains were collected, preserved, and processed for analysis. Retinal cross-sections from various geometric subregions and brain cross-sections from the frontal cortex (A-9 area) were prepared for histological analyses through immunostaining and silver staining. Information on the antibodies that were used is provided in **Supplementary Table 2**.

### 2.4 Microscopy and histological quantification in pre-defined geometrical regions

Systematic quantitation of retinal NFTs, oligo-tau, p-tau, PHF-tau, and Cit-tau in NIH Image J was performed on microscopic images captured using fluorescence and a bright-field Carl Zeiss Axio-Imager Z1 microscope with ZEN 2.6 blue edition software; a stereological method previously described ^24^. On average, 12 images (20× magnification) were obtained from each patient for quantification (see details in Supplementary Methods).

### 2.5 GeoMx digital spatial profiling for tau proteins

Formalin fixed paraffin embedded human brain and retinal cross-sections were prepared for a p-tau panel and analyzed by GeoMx® digital spatial profiler (DSP) Proteomics (see details in Supplementary Methods).

### 2.6 Statistical Analysis

GraphPad Prism Software version 8.3.0 was used for analyses. One-way ANOVA followed by Tukey’s multiple comparison post-test was used to analyze statistical significance between groups. Pearson’s correlation coefficient (*r*) test was used to determine associations between markers in this study.

## 3. Results

To investigate the burden and spatiotemporal distribution of abnormal tau forms (**Supplementary Figure 1a**) in the retina, we prepared retinal cross-sections from the superior- and inferior-temporal (ST/IT) regions in a cohort of patients with premortem diagnosis of MCI (due to AD; n=11, mean age 89.6 ± 6.1 years, 7 females/4 males) or AD dementia (n=34, mean age 83.9 ± 12.7 years, 21 females/13 males) and postmortem neuropathologically confirmed as AD, and individuals with NC (n=30, mean age 81.5 ± 10.0 years, 17 females/13 males). No significant differences were found in age, sex, or post-mortem interval (PMI) among the three diagnostic groups. Demographic information, brain neuropathology, and retinal tauopathy histological biomarkers are detailed in **Table 1**. The correlations of retinal tau biomarkers with brain neuropathology and cognitive scores are summarized in **Table 2**.

**Table 2.** Correlation analysis between retinal tauopathy markers and brain neuropathology and cognitive decline.

|  | Brain | Core AD pathology |  | Non-specific tissue injury | Vascular co-morbidity | AD staging |  | Cognition |
| --- | --- | --- | --- | --- | --- | --- | --- | --- |
| Retinal Tauopathy (Antibody) | | Aβ-P | NFT | Atrophy | CAA | Braak | ABC | CDR |
| pS202/T205-tau (AT8) | *r* [*n*] | 0.15 [33] | 0.2 [33] | -0.07 [33] | **0.36** [33] | 0.17 [33] | 0.23 [33] | **0.55** [29] |
|  | *P* | n.s. | n.s. | n.s. | **0.0399 \*** | n.s. | n.s. | **0.0018 \*\*** |
| pS212/T214-tau (AT100) | *r* [*n*] | **0.61** [13] | **0.66** [13] | 0.25 [13] | -0.12 [13] | 0.47 [13] | -0.13 [8] | -0.23 [9] |
|  | *P* | **0.0266 \*** | **0.0145 \*** | n.s. | n.s. | 0.1058 | n.s. | n.s. |
| pS396-tau (pS396) | *r* [*n*] | **0.50** [37] | **0.68** [37] | 0.13 [37] | 0.13 [34] | **0.44** [37] | **0.52** [37] | **0.37** [32] |
|  | *P* | **0.0017 \*\*** | **<0.0001 \*\*\*\*** | n.s. | n.s. | **0.0061 \*\*** | **0.0010 \*\*** | **0.0398 \*** |
| CitR<sub>209</sub>-tau | *r* [*n*] | 0.25 [33] | 0.17 [33] | 0.04 [33] | **0.35** [33] | 0.2 [33] | 0.33 [33] | **0.53** [29] |
|  | *P* | n.s. | n.s. | n.s. | **0.0475 \*** | n.s. | 0.0596 | **0.0030 \*\*** |
| Oligo-tau (T22) | *r* [*n*] | **0.41** [34] | **0.53** [34] | 0.02 [34] | **0.60** [32] | **0.60** [34] | **0.41** [32] | **0.46** [27] |
|  | *P* | **0.0168 \*** | **0.0012 \*\*** | n.s. | **0.0003 \*\*\*** | **0.0002 \*\*\*** | **0.0197 \*** | **0.0150 \*** |
| PHF-tau | *r* [*n*] | 0.32 [18] | **0.50** [18] | 0.43 [18] | 0.48 [16] | **0.52** [18] | **0.64** [18] | 0.16 [14] |
|  | *P* | 0.1925 | **0.0335 \*** | 0.0746 | 0.0623 | **0.0264 \*** | **0.0043 \*\*** | 0.5738 |
| NFTs (MC-1) | *r* [*n*] | -0.07 [36] | 0.17 [36] | **0.41** [36] | **0.35** [33] | 0.16 [36] | 0.11 [34] | 0.17 [28] |
|  | *P* | n.s. | n.s. | **0.0129 \*** | **0.0427 \*** | n.s. | n.s. | n.s. |

Pearson's coefficient (*r*) correlation determining the strength of association between retinal tauopathy markers vs. brain neuropathology and cognitive decline. *P* and *r*-values are shown. *P* and *r*-values presented in bold type demonstrate significance. Antibody name is listed following each retinal tauopathy marker. Mean ABC scores were determined as follows: A, Aβ plaque score modified from Thal; B, NFT stage modified from Braak; C, neuritic plaque score modified from CERAD. Abbreviations: Aβ , amyloid beta-protein; AD, Alzheimer's disease; CAA, cerebral amyloid angiopathy; CDR, Clinical Dementia Rating; MMSE, Mini-Mental State Examination; NFTs, neurofibrillary tangles; n.s. not significant.

### 3.1. Identification and quantification of NFTs and oligomeric tau in the MCI and AD retina

In AD brains, NFTs represent mature, aggregated, and late-stage forms of p-tau buildup. Retinal NFTs were identified and quantified in a sub-cohort of donors with MCI (n=9, mean age 89.7 ± 5.1 years, 5 females/4 males), AD (n=21, mean age 85.6 ± 8.5 years, 11 females/10 males), and NC controls (n=19, mean age 84.5 ± 9.1 years, 9 females/10 males), and were compared to brain NFTs. Immunohistochemical (IHC) staining for conformational- and sequence-specific NFTs was conducted using the MC-1 monoclonal antibody, which identifies the aa 7-9 and 312-322 tau epitopes ^34^. MC-1 primarily recognizes pre-tangles and mature tangles ^35^. We found occasional retinal MC-1^+^ NFTs in the ganglion cell layer (**Figure 1a**, upper panel) and co-staining of MC-1^+^ with βIII-tubulin cell within the INL of MCI and AD patients compared to NC controls (**Figure 1a**, lower panels). The structure of retinal NFTs, as revealed by MC-1 and Bielschowsky silver staining, appears to resemble those in the brain (**Figure 1a-1b**; additional Bielschowsky silver stain images across retinal layers and in paired brain tissues from other AD patients in **Supplementary Figure 1**). Quantitative analysis of brain NFTs measured by Gallyas Silver and Thioflavin staining was 3.2-fold higher in MCI and 4.1-fold higher in AD, compared to NC controls. Stereological quantification of MC-1-positive NFTs in the respective retinas indicates significant and modest 1.8-fold and 2.0-fold increases in MCI and AD patients compared to NC controls, respectively (**Figure 1b**). Indeed, there was a considerable amount of overlap in the levels of retinal MC-1 percent area in older individuals with normal cognition and those with MCI due to AD and AD dementia. Pearson’s correlation coefficient (*r*) analysis demonstrates that retinal MC-1-positive NFTs burden moderately and significantly associates with the severity of brain atrophy (**Figure 1b** and **Table 2**) and cerebral amyloid angiopathy (CAA, **Table 2**).

**Figure 1.**
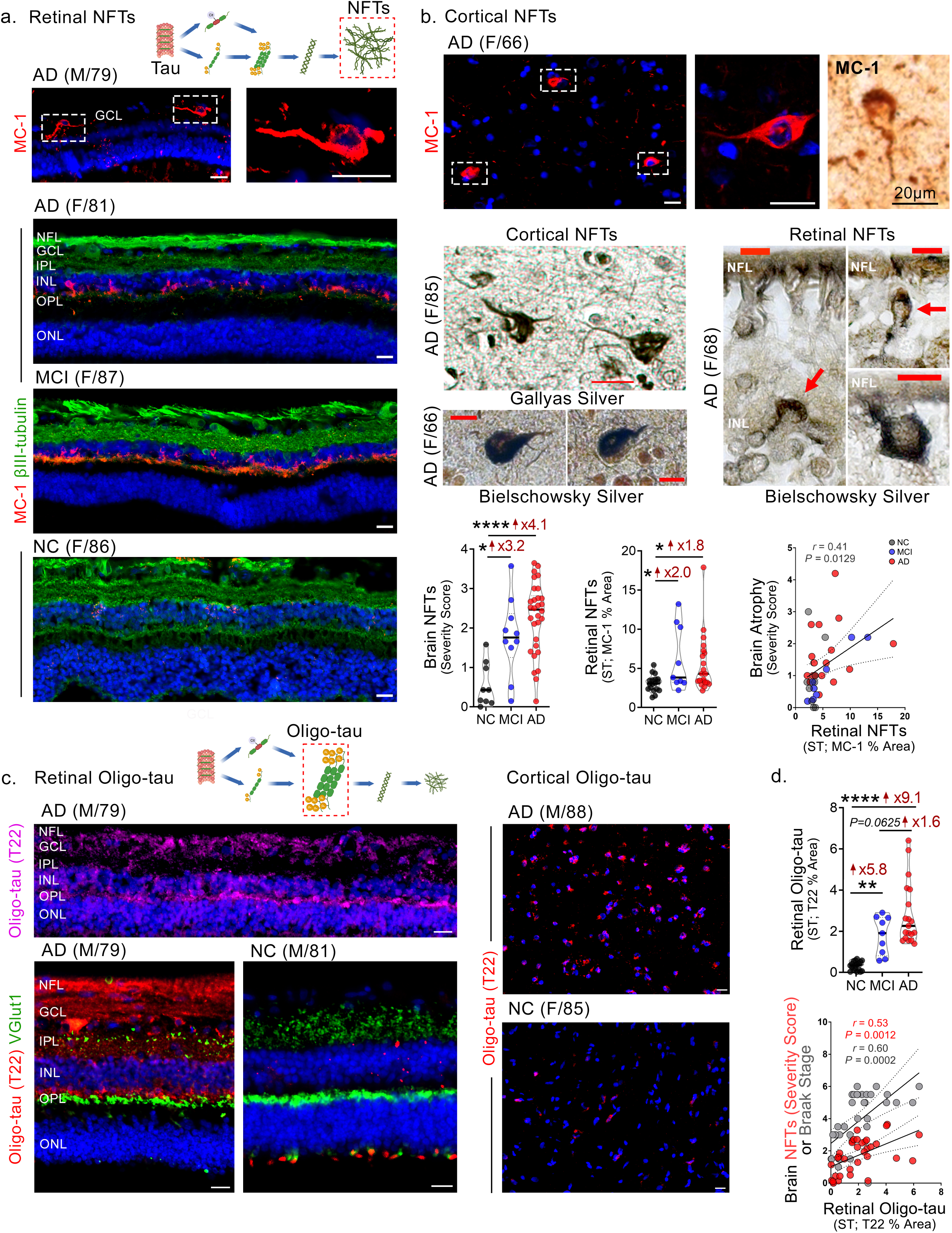
Identification of neurofibrillary tangles and oligomeric tau in the retina of MCI and AD patients. **a.** Representative images of immunofluorescent staining for neurofibrillary tangles (NFTs, MC-1, red), βIII-tubulin (green), and DAPI (blue) on postmortem cross-sections of retina from normal cognition (NC, n=19 [control]) patients as well as from patients with mild cognitive impairment (MCI, n=9) and those with Alzheimer’s disease (AD, n=21). All scale bars=20µm. **b.** Representative images of cortical NFTs stained by immunofluorescence (red) and peroxidase-based DAB (brown) with the MC-1 antibody (Scale bars=20µm), Gallyas silver (Scale bars=20µm), and Bielschowsky silver (Scale bars=10µm) staining on brain and retinal sections. Bar graph of brain NFT severity scores. Quantitative analysis of retinal NFTs immunoreactivity (IR) (n=49 in total). Pearson’s correlation coefficient (*r*) between brain atrophy severity scores and retinal NFTs burden (n=36). **c.** Representative images of immunofluorescent staining for oligomeric tau (oligo-tau, T22, magenta or red), vesicular glutamate transporter 1 (VGlut1, green), and DAPI (blue) on postmortem cross-sections of retina from normal cognition (n=19) patients as well as from patients with MCI (n=9) and those with AD (n=21). All scale bars=20µm. **d.** Quantitative analysis of retinal T22^+^oligo-tau IR (n=49 in total). Pearson’s correlation coefficient (r) between retinal T22^+^oligo-tau and brain NFTs severity score or Braak stage of brain NFTs (n=34). Data from individual donors (circles) as well as group means ± SDs are shown. \**p* < 0.05, \*\**p* < 0.01, \*\*\*\**p* < 0.0001, by one-way ANOVA with Tukey’s post-hoc multiple comparison test. Fold changes are shown in red. Abbreviations: F, female; M, male; age (in years). Nerve fiber layer (NFL), ganglion cell layer (GCL), inner plexiform layer (IPL), inner nuclear layer (INL), outer plexiform layer (OPL), and outer nuclear layer (ONL). Illustrations created with Biorender.com.

We then explored the presence of toxic, propagating, oligomeric tau forms in the retina of MCI and AD patients. In AD brains, oligo-tau forms are assembled from small p-tau aggregates after dislodging from microtubules in neurons ^2^. Previous research demonstrated that extracting oligo-tau from the AD brain using the T22 antibody and injecting these oligomers into wild-type mouse brains caused neurotoxicity and the propagation of abnormal endogenous murine tau ^36^. Here, we stained ST/IT retinal and brain cross-sections, from the same cohort as described above, for T22-positive oligo-tau, alone and in combination with the pre-synaptic marker, vesicular glutamate transporter 1 (VGlut1; **Figure 1c**). We observed abundant cellular and diffused oligo-tau, especially in the synaptic-rich layers (OPL, IPL), with scarce VGlut1^+^ signals in the retinas of AD patients compared to NC controls (**Figure 1c**, left panel). The differences in brain T22-positive oligo-tau between AD and NC retinas are also shown (**Figure 1c**, right panel). Stereological analysis demonstrated a substantial 9.1-fold increase in retinal oligo-tau in AD patients and a 5.8-fold increase in MCI patients compared to NC controls (**Figure 1d**, upper graph). Notably, Pearson’s correlation analyses revealed that retinal T22-positive oligo-tau moderately correlates with brain NFT burden and strongly with the Braak staging, a parameter of tauopathy spread across brain regions during AD progression (**Figure 1d**, lower graph). Furthermore, retinal oligo-tau correlated strongly with CAA severity, and moderately with brain Aβ burden, A(amyloid-beta plaque) B(NFT stage) C(Neuritic plaque)—ABC score, and the clinical dementia rating (CDR) cognitive score (**Table 2**).

### 3.2. GeoMx tauopathy profiling and p-tau histopathology in the retina of MCI and AD patients

We employed the high-throughput NanoString GeoMx® digital spatial profiling (DSP) technique (**Figure 2a)** to determine protein quantities for total tau and different p-tau forms in retinal cross-sections and paired brain tissues prepared from a donor cohort consisting of MCI (n=6, mean age 88.5 ± 5.0 years, 3 females/3 males), AD (n=9, mean age 85.1 ± 7.8 years, 5 females/4 males), and NC controls (n=9, mean age 89.3 ± 9.4 years, 6 females/3 males). We detected higher total tau levels in the AD brain and MCI/AD retina compared to NC controls (**Figure 2a**; bottom left graph). The GeoMx p-tau module included an analysis of phosphorylation sites at serine 199 (S199), serine 214 (S214), serine 396 (S396), serine 404 (S404), and threonine 231 (T231). Significant increases in retinal S214, S396, S404, and T231-p-tau were detected in MCI patients compared to NC controls, and retinal pS396-tau was also significantly elevated in AD patients (**Figure 2a**; lower panel). In a subset of paired brains, significant increases in brain pS214-, pS396-, and pS404-tau were found in AD patients. Interestingly, higher levels of retinal pT231- and pS404-tau were noted in MCI versus AD patients, suggesting early retinal tau changes during AD progression (**Figure 2a**). Our quantitative GeoMx analysis in this cohort indicated that retinal pT231-tau significantly correlates with brain NFTs and neuropil threads (NT) burdens (**Figure 2a**, right), along with Aβ burden and CAA scores (**Supplementary Figure 2a-b**). Furthermore, retinal pS214-tau strongly correlated with cognitive scores assessed by mini-mental state examination (MMSE) (**Supplementary Figure 1c**).

**Figure 2.**
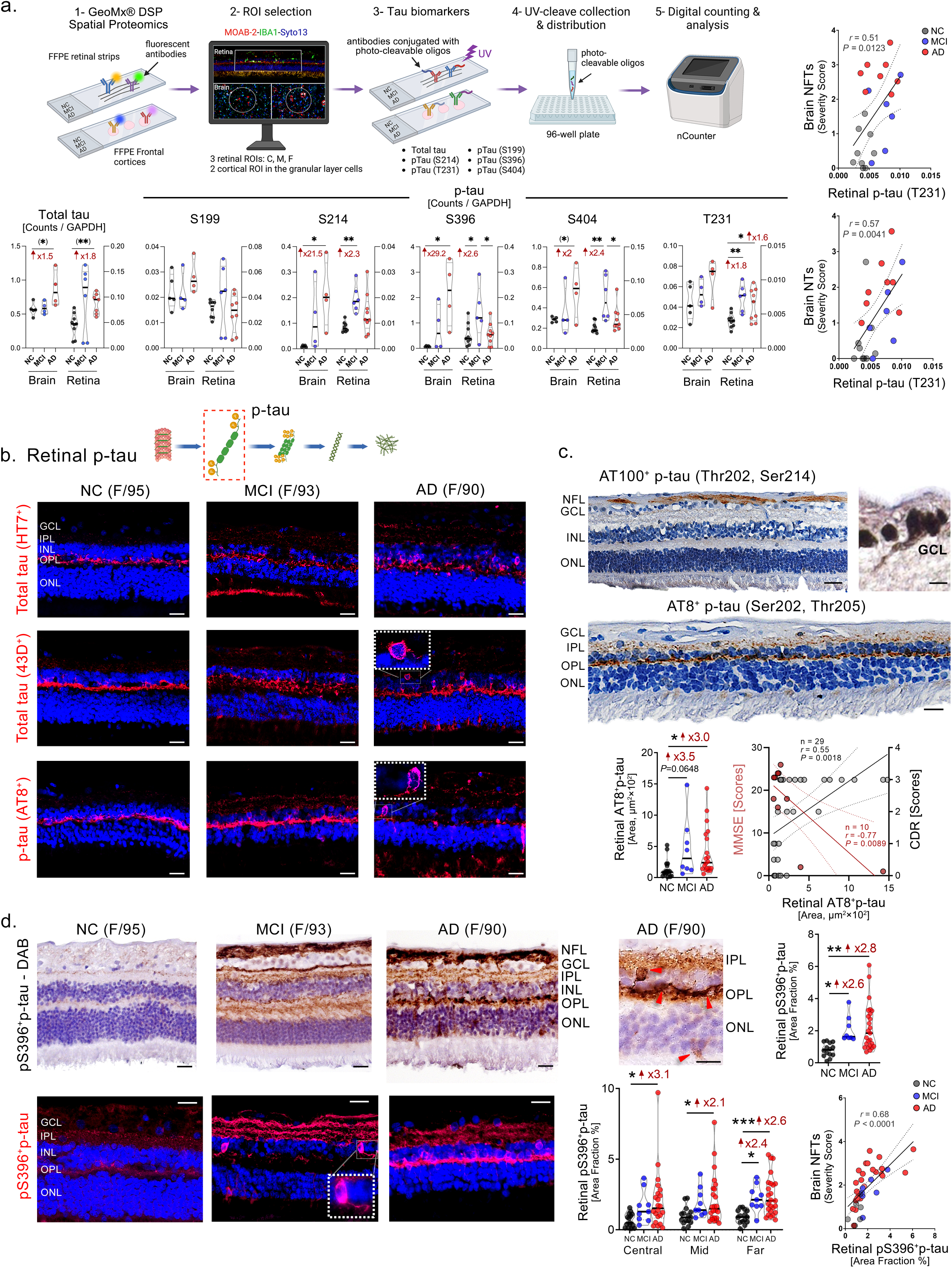
Retinal tau and phosphorylated-tau forms in MCI and AD patients determined by spatial profiling and histological examination. **a.** Graphical illustration of NanoString GeoMx® DSP analyses for tau proteins in retinal and respective brain samples. Quantitative analysis of retinal total tau and p-tau at phosphorylation sites of S199, S214, S396, S404, and T231 detected by GeoMx® in retinas from AD (n=9) and MCI (n=6) patients, and normal cognition controls (NC; n=9), and a subset of paired-brain tissues (A-9, frontal cortex region; n=13). Pearson’s correlation coefficient (*r*) analyses are shown between retinal pT231-tau and the severity of brain neurofibrillary tangles (NFTs, n=23) or neuropil threads (NTs, n=23). **b.** Representative images of immunofluorescent staining for total tau (HT7^+^ or 43D^+^) and AT8^+^p-tau (S202/T205) in retinal cross-sections from human donors with NC, MCI, and AD. Scale bars=20µm. **c.** Representative images of peroxidase-based DAB staining of AT100^+^p-tau (T202/S214) and AT8^+^p-tau (S202/T205) on retinal cross-sections from AD patients. Quantitative analysis of retinal AT8^+^p-tau immunoreactivity (n=46 in total: n=20 AD, n=8 MCI, n=18 NC). Pearson’s correlation coefficient (*r*) analysis between retinal AT8^+^p-tau and the cognitive scores as assessed by mini-mental state examination (MMSE, n=10) or Clinical Dementia Rating (CDR, n=29). **d.** Representative images of immunofluorescent and peroxidase-based staining of pS396-tau in retinal cross-sections from AD (n=25) and MCI (n=8) patients, and NC controls (n=14). Scale bars=20µm. Quantitative analysis of pS396-tau immunoreactive area in ST/IT retina and in separate central-, mid-, and far-peripheral retina subregions (n=47 in total). Pearson’s correlation coefficient (*r*) analysis between retinal pS396-tau versus respective brain NFTs burden (n=37). Data from individual donors (circles) as well as group means ± SDs are shown. \**p* < 0.05, \*\**p* < 0.01, \*\*\**p* < 0.001, by one-way ANOVA with Tukey’s post-hoc multiple comparison test for group analyses. Two group comparisons by student t-test are indicated in parenthesis. Fold changes are shown in red. Abbreviations: F, female; M, male; age (in years). Nerve fiber layer (NFL), ganglion cell layer (GCL), inner plexiform layer (IPL), inner nuclear layer (INL), outer plexiform layer (OPL), and outer nuclear layer (ONL). Illustrations created with Biorender.com.

The detection of significant changes in retinal and brain p-tau forms in GeoMx DSP analysis prompted an additional histological investigation of other retinal p-tau epitopes, including pS396-tau, in a larger cohort. IHC analysis of total tau using both HT7 and 43D antibodies revealed a considerable retinal tau signal, mostly restricted to the outer plexiform layer (OPL) in NC subjects, spreading across all retinal layers in MCI and AD patients (**Figure 2b**). In a subset of donors, the analysis of retinal pT202/S214-tau (using the AT100 antibody) showed a non-significant trend of increases in MCI and AD patients compared to NC controls (**Figure 2c, Supplementary Figure 3a-b**). Nevertheless, retinal AT100^+^p-tau deposition was strongly associated with both brain Aβ and NFT burdens (**Supplementary Figure 3c**). Examination of retinal AT8-positive signals of p-tau at S202/T205 epitopes detected a 3.5-fold trend of increase in MCI and a significant 3-fold increase in AD, compared to NC controls (**Figure 2b** and **2c**, lower panels). Retinal AT8-positive p-tau deposition weakly associated with CAA severity scores (**Table 2**) but showed a moderate-to-strong correlation with cognitive scores, as determined by CDR and MMSE (**Figure 2c**).

Analysis of retinal pS396-positive tau in a larger cohort, using immunofluorescence and peroxidase-based immunostaining, shows substantial pS396-tau accumulation across all retinal layers in MCI and AD patients compared to NC (**Figure 2d**; intraneuronal p-tau structures are often detected). In agreement with the quantitative GeoMx DSP findings (**Figure 2a**), stereological analysis of pS396-tau revealed an early and significant 2.6-fold increase in MCI and a 2.8-fold increase in AD compared to NC retinas (**Figure 2d**, upper right). Examination of retinal pS396^+^p-tau distribution per retinal region indicates that the far-peripheral retina exhibits an earlier and more significant increase of these p-tau forms, providing clearer separation between the diagnostic groups (**Figure 2d**; representative images per retinal C/M/F regions in **Supplementary Figure 4**). A strong and highly significant correlation was identified between retinal pS396^+^p-tau burden and brain NFTs severity scores (**Figure 2d**). Notably, retinal pS396^+^p-tau deposition also significantly associated with brain Aβ burden, ABC scores, Braak staging, and CDR cognitive scores (**Table 2**).

### 3.3. Early tau citrullination and late PHF-tau increases in the AD retina

Upon detecting increased levels of oligo-tau, p-tau, and mature NFT forms in the retina of MCI and AD patients, we further examined the pre-NFT forms—the PHF-tau aggregates—in the AD retina. Previously, brain PHF-tau in AD has been associated with chronic neuroinflammation, including activated microgliosis ^37,38^. Additionally, tau-laden neurons were susceptible to excessive microglial synaptic pruning ^38^. Retinal IHC analysis, using the PHF-1 antibody recognizing pS396- and pS404-tau in paired helical filaments, was performed on a subset of patient donors with MCI (n=5, mean age 89.8 ± 5.8 years, 3 females/2 males), AD (n=10, mean age 88.1 ± 7.4 years, 5 females/5 males), and NC controls (n=9, mean age 82.5 ± 8.5 years, 3 females/5 males). This analysis showed marked PHF-tau accumulations across retinal layers in AD patients, particularly localized in VGLUT1^+^ synaptic-rich and IBA1^+^ microgliosis regions (**Figure 3a**, left panel). Quantitative IHC analysis revealed a significant 2.3-fold increase in retinal PHF-tau in AD, but not in MCI, compared to NC controls (**Figure 3a**, right upper). Retinal PHF-tau deposition strongly associated with brain NT burden and ABC scores (**Figure 3a**, right lower), and moderately with brain NFTs burden and Braak staging (**Table 2**).

**Figure 3.**
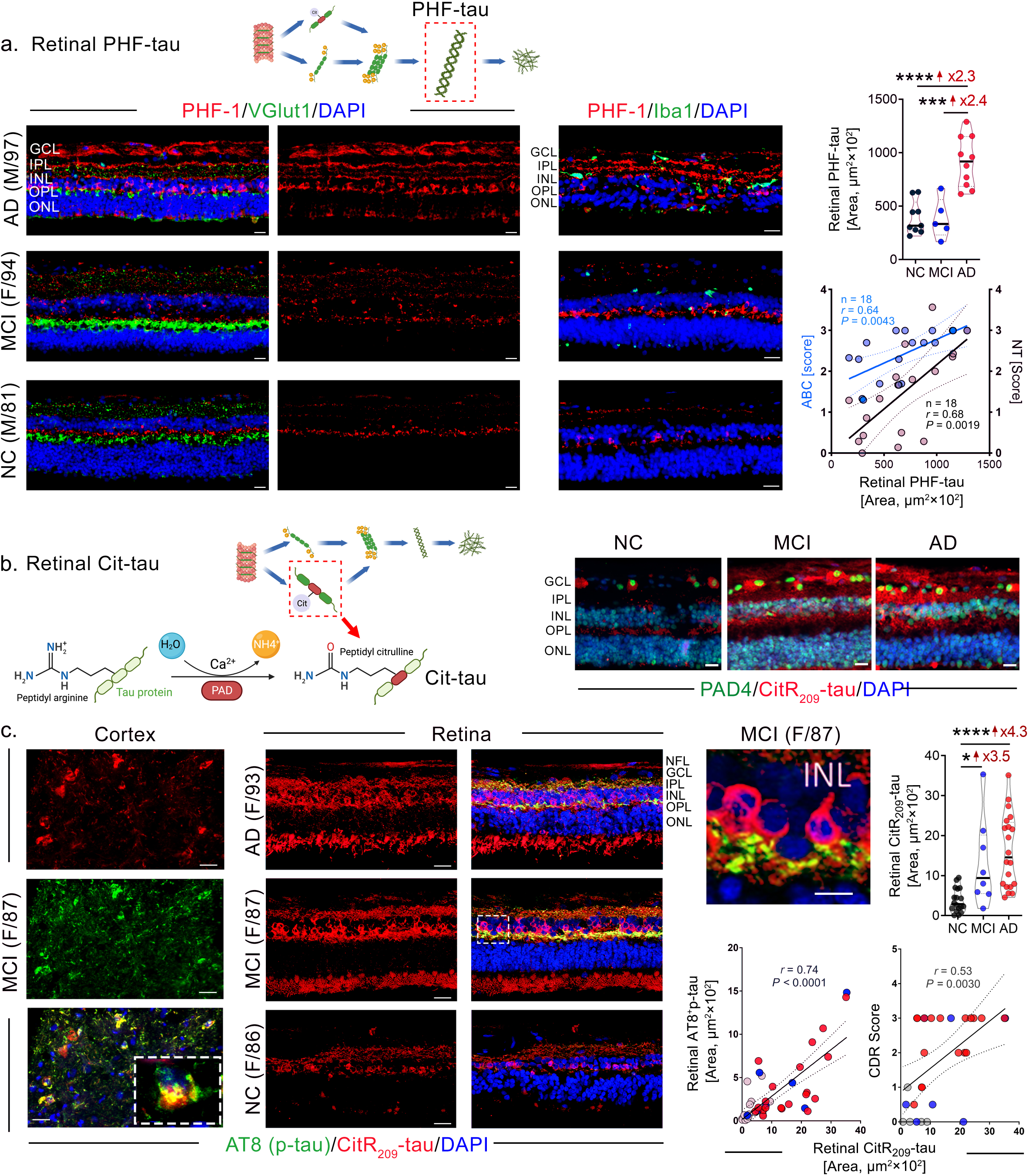
Accumulation of paired-helical filaments and citrullinated-tau forms in the AD retina. **a.** Representative images of immunofluorescent staining for PHF-1^+^ paired-helical filaments of tau (PHF-tau, red), vesicular glutamate transporter 1 (VGlut1) or IBA1 (green), and DAPI (blue) in retinal cross-sections from patients with MCI (n=5) and AD (n=10), and NC controls (n=9). Scale bars=20µm. Quantitative analysis of retinal PHF-tau immunoreactive area (n=24 in total). Pearson’s correlation coefficient (*r*) analyses of retinal PHF-tau against brain neuropil threads (NT, n=18) or A(amyloid-beta)/B(Braak stage)/C(CERAD) average scores (n=18). **b.** Graphical illustration of tau citrullination post-translation modification catalyzed by protein arginine deiminase (PAD) enzymes. Representative images of immunofluorescent staining for PAD4 enzyme (green), citrullinated tau at arginine R209 site (CitR_209_-tau, red), and DAPI (blue) in retinal cross-sections from MCI and AD patients as compared with NC controls. Scale bars=20µm. **c.** Representative images of immunofluorescent staining for AT8^+^p-tau (green), CitR_209_-tau (red), and DAPI (blue) in brain and retinal cross-sections from patients with AD (n=20), MCI (n=8), and NC controls (n=18). Scale bars=20µm. Quantitative analysis of CitR_209_-tau immunoreactive area (n=46 in total). Pearson’s correlation coefficient (*r*) analyses between retinal CitR_209_-tau and retinal AT8^+^p-tau (n=46) or cognitive scores by Clinical Dementia Rating (CDR, n=29). Data from individual donors (circles) as well as group means ± SDs are shown. \**p* < 0.05, \*\*\**p* < 0.001, \*\*\*\**p* < 0.0001, by one-way ANOVA with Tukey’s post-hoc multiple comparison test for group analyses. Fold changes are shown in red. Abbreviations: F, female; M, male; age (in years). Nerve fiber layer (NFL), ganglion cell layer (GCL), inner plexiform layer (IPL), inner nuclear layer (INL), outer plexiform layer (OPL), and outer nuclear layer (ONL). Illustrations created with Biorender.com.

Citrullination is a post-translational modification in which an arginine amino acid is converted to a citrulline amino acid, a process catalyzed by peptidyl arginine deiminase (PAD) enzymes that play a significant role in several chronic diseases ^39^. Neurons expressing PAD4 were found to accumulate citrullinated proteins in AD cortices and hippocampi ^40^. A recent study identified altered PAD4 expression and cit-tau accumulation in the brains of AD patients ^4^. Aberrant tau deposition activated PAD4 in neurons, which also led to citrullination of tau at multiple arginine residues ^4^. In this study, we detected prominent neuronal PAD4 expression along with marked accumulations of retinal and cortical citrullinated arginine (R)-209 tau (CitR_209_-tau) and AT8^+^p-tau (S202/T205) in MCI and AD patients compared to NC controls (**Figure 3b-c**). In both retinal and paired brain tissues, we found increased co-localized signals of AT8^+^p-tau and CitR_209_-tau in MCI and AD patients, with clear intra-neuronal CitR_209_-tau signals in the inner nuclear layer – INL (**Figure 3c**; higher magnification of MCI retina). Moreover, Pearson’s correlation analysis indicated a strong positive association between these two post-translational modifications of tau (CitR209tau and AT8^+^p-tau) in the retina (**Figure 3c**, lower right panel). Stereological quantification of retinal CitR_209_-tau indicates a substantial 3.5-fold and 4.3-fold increase in MCI and AD patients, respectively, compared to NC controls (**Figure 3c**). Retinal CitR_209_-tau moderately correlated with CDR cognitive scores (**Figure 3c**) and weakly with CAA scores (**Table 2**). Analyses of retinal AT8^+^p-tau, PHF-tau, and CitR_209_-tau spatial distribution per sub-regions indicated that while both AT8^+^p-tau and PHF-tau increase earlier and/or more significantly in the mid- and far-peripheral retina, CitR_209_-tau appeared earlier and more pronouncedly in the central retina (**Supplementary Figure 5**).

## 4. Discussion

In this study, we present the first evidence of retinal oligomeric tau and citrullinated tau buildup, alongside the identification of classical structures of NFTs in the retina of MCI and AD patients. Moreover, we find that early-stage hyper-phosphorylated and citrullinated tau, propagating-type tau oligomers, and mature NFT forms accumulate in the retinas of patients with early AD (MCI due to AD) and late AD dementia stages, while retinal PHF-tau increases only in the later stages of AD progression. GeoMx spatial profiling analyses reveal site-specific increases in various tau phosphorylation epitopes in the AD retina, particularly at the MCI stage. Importantly, retinal tau oligomers exhibit the largest magnitude of increases in MCI and further in AD patients compared to matched NC controls, showing the strongest correlation with Braak staging, which represents the spread of brain tauopathy. Overall, our data demonstrate that most retinal tauopathy forms are increased in early AD and correlate with one or more AD neuropathology and cognitive parameters, guiding the development of future retinal tau biomarkers for AD screening and monitoring.

Aggregates of hyperphosphorylated tau in the form of brain NFTs are core hallmarks that define AD diagnosis and are tightly linked to neurodegeneration ^34,41^. In agreement with this, our study identifies NFTs in the MCI and AD retina, the only abnormal tau form significantly correlated with the severity of brain atrophy. Despite some contradictions, several previous reports have found an association between neocortical NFT burden and antemortem cognitive decline in AD cases ^42-47^. In our cohort, there were modest increases in retinal MC-1-positive NFTs in MCI and AD patients, with no association between retinal NFTs burden and cognitive status, as assessed by CDR scores. This suggests that mature fibrillary tau forms may not affect the retina to the same extent as the brain during AD progression. Previous studies have described the accumulation of diverse p-tau forms and NFT-like structures in postmortem retinal tissues from AD patients ^18-23,25,26,28,30-32,48^. A recent study reported a mild immunohistochemical NFT signal in the retinal outer plexiform layer (OPL), inner nuclear layer (INL), and inner plexiform layer (IPL) from AD cases using the MC-1 antibody, which specifically recognizes conformational NFTs ^32^. However, none have successfully detected the typical patterns of mature fibrillary tau forms such as PHF-tau or NFTs in the retina. In our study, immunohistochemical staining using the PHF-1 and MC-1 antibodies, and the Bielschowsky silver stain, validated the presence of typical paired-helical filaments of tau and NFTs in the retina of MCI and AD patients. Retinal PHF-tau and NFTs appear structurally similar to their cerebral counterparts and are frequently found within neurons located in the inner nuclear and ganglion cell layers. The inconsistency in previous observations is likely due to differences in tissue preservation, fixation, staining techniques, and the analyzed geometric regions ^17^.

Oligomers of tau are considered highly neurotoxic intermediate assemblies that are precursors of protofilaments, PHF-tau, and subsequent NFTs ^2^. In the AD brains, tau oligomers spread across anatomical regions and are involved in the early stages of AD pathogenesis ^2,49^. These oligomers have been shown to impair neuronal energy production, synaptic integrity, microtubule assembly, and axonal transport ^50^. In this study, we identified a substantial accumulation of T22-positive tau oligomers in the retina of MCI and AD patients compared to normal controls. Both diffused and intracellular tau oligomer signals were observed across the retinal layers, predominantly spanning from the OPL through the nerve fiber layer (NFL). Among the diverse retinal tau forms measured in this study, tau oligomers exhibited the earliest and most extensive increases in the AD retinas, showing strong associations with Braak staging and CAA severity scores. Therefore, tau oligomers should be evaluated as a potential retinal tau marker for early detection and tracking of AD progression in future studies.

Previous studies utilizing immunohistochemistry and western blot on postmortem AD donor retinas have identified retinal p-tau at phosphorylation sites, including S202, T205, T217, T212, S214, T181, T231, S396, and S404 ^18-21,31,32^. In this study, we employed the GeoMx spatial profiling method and detected increased total tau in AD brains and retinas, aligning with previous findings in the brain ^51^. Quantitative GeoMx analysis also indicated increases in retinal p-tau at S214, S396, S404, and T231 sites in MCI patients compared to controls, with moderate to strong associations between retinal pT231-tau and brain neurofibrillary tangles and neuropil threads. Interestingly, retinal pT231- and pS404-tau levels were significantly higher in MCI patients compared to AD dementia patients, suggesting their earlier accumulation during AD progression. This aligns with a previous study showing increased pT231-tau levels in the CSF of preclinical AD patients ^52^. Histological examinations indicated strong correlations between additional retinal p-tau types and brain pathology or cognitive status. Retinal AT8-positive pS202/T205-tau burden strongly correlated with MMSE cognitive scores, whereas pS396-tau and AT100-positive pS212/T214-tau forms strongly associated with brain NFT burden. Of note, a previous study reported that retinal pS202/T205-tau correlated with brain NFT levels in the hippocampus, temporal pole, medial frontal gyrus, and the parietal lobe ^21^. Therefore, retinal pS202/T205-tau and pS396-tau forms should be further considered as potential markers to track brain NFTs severity and cognitive decline. Collectively, these results suggest that retinal p-tau accumulation may occur early in AD pathogenesis and may predict brain tauopathy and cognitive status.

In a previous study, intense accumulation of retinal PHF-tau in the IPL, OPL, and INL of AD patients was reported ^32^. We observed that PHF-tau is not restricted to these three layers but also found it extends to the retinal nerve fiber and ganglion cell layers (NFL, GCL) in AD patients. The abnormal folding of hyperphosphorylated tau at both S396 and S404 sites leads to the generation of insoluble PHF-tau ^53^. While GeoMx data show early upregulation of each pS396- or pS404-tau forms in MCI retinas, histological analysis of conformational PHF-tau structures indicated increases later in the AD retina compared to the control retina. This result indicates that, unlike early hyperphosphorylation of tau, its folding into PHF tau may occur in the retina at later disease stages. Correlation analyses further suggest that retinal PHF tau may be a strong predictor to track brain ABC, NFTs, and NTs scores in patients.

The current study provides the first evidence of hyper-citrullinated tau in the human MCI and AD retina. Post-translational modifications of tau are a prerequisite for the formation of PHF-tau and NFTs ^54^. Tau is a client of the PAD4 enzyme, which can cause irreversible citrullination of its arginine residues ^4^. In our study, we identified early increases in retinal CitR_209_-tau in MCI and AD patients. In the OPL and INL, we observed a considerable amount of co-localization and close interplay between retinal CitR_209_-tau and AT8-positive pS202/T205-tau signals, suggesting that citrullination and hyperphosphorylation both occur during the development of retinal tauopathy in AD. Furthermore, similar to retinal AT8-positive pS202/T205-tau, retinal CitR_209_-tau also exhibited significant correlations with cognitive status. Since hyper-citrullination of proteins has been implicated in multiple chronic human diseases ^39,55^ and citrullinated tau may impact oligomerization and microglial activation ^4^, our results of increased retinal CitR_209_-tau perhaps imply altered aggregation, propagation, and clearance properties of tau in the MCI and AD retina.

Despite the many strengths of this study in providing insights into various aberrant tau forms associated with AD in the retinas of both MCI and AD patients and defining their relationship to disease status, we acknowledge a few limitations. This is a cross-sectional case-control study primarily focused on correlation; therefore, caution must be exercised before implicating cause-and-effect conclusions. In addition, this is the first study to analyze tauopathy in rare retinal tissues from MCI (due to AD) patients. However, we recognize that a larger sample size is warranted in future studies. Given the inconsistencies in previous reports on retinal AD tauopathy—where two recent studies identified certain pathological tau forms in the AD retina ^21,32^ that previous reports could not detect ^56,57^ or only partially detected ^18,31^—workshop groups focused on harmonizing the methodologies for analyzing retinal tauopathy will be useful in future studies.

In summary, our study provides evidence of diverse and novel pathological tau forms in the retina of MCI and AD patients. These include classical conformational tau aggregates such as paired helical filaments and neurofibrillary tangles, tau oligomers, and hyper-phosphorylated and citrullinated tau forms. Our findings suggest that the retina is highly affected by tauopathy during AD pathogenesis and that aberrant retinal tau forms can serve as predictors of brain pathology and cognitive decline. Taken together, this study offers valuable insights into AD-related retinal tau forms, which have the potential to be utilized in the future development of noninvasive and high spatial-resolution retinal tau imaging tools for the early detection and monitoring of AD.

## Acknowledgments

We thank Elijiah Maxfield for assisting with manuscript editing. We thank the Cedars-Sinai Medical Center Pathology and Imaging Core for assistance with GeoMx experiment. We acknowledge the contribution of Prof. Carol Ann Miller, the former director of the USC-ADRC neuropathology laboratory, for providing part of the neuropathological reports. The authors dedicate this manuscript to the memory of Dr. Salomon Moni Hamaoui and Lillian Jones Black, both of whom died from Alzheimer’s disease.

## Consent statement

This study is not considered a human subjects research and we confirm that consent was not necessary, for the reasons described as follow: we processed and analyzed deidentified retinal tissues of deceased patients that were provided by the USC-ADRC (C.A.M, D.H.) and from the NDRI (A.V.L.).

## Author contributions

H.S., N.M., Y.K., M.K.H.: designed and performed experiments, collected and analyzed data, created figures, drafted, and edited the manuscript. D-T.F., M.R.D.: performed experiments, collected and analyzed data, and made the illustrations. E.R., G.M.B., O.J., A.R.: performed experiments and analyses. V.R., W.G.T.: supervised the NanoString GeoMx experiment. R.K., M-L.B.S., J.F.B., D.C.L.: provided antibodies and advised regarding the oligo-tau, PAD4, and Cit-tau analyses. A.V.L., A.A.K., L.S.S., J.A.S., D.H.: provided donor eyes and clinical reports and edited the manuscript. K.L.B. assisted with data interpretation and manuscript editing. M.K.H. was responsible for study conception and design, data analysis and collection, interpretation of data, study supervision, and manuscript writing and editing. All authors have read and approved of this manuscript.

