## Supplementary materials for "Identification of retinal tau oligomers, citrullinated tau, and other tau isoforms in early and advanced AD and relations to disease status"

**eMethods.** Extended method information.

**eTable 1.** List of human donors in this study.

**eTable 2.** List of antibodies.

**eFigure 1.** Bielschowsky silver staining for brain and retinal NFTs.

**eFigure 2.** Additional correlations between GeoMx® DSP of retinal p-tau and brain pathology.

**eFigure 3.** Retinal AT100$^{+}$p-tau burden and association with brain pathology.

**eFigure 4.** Regional difference of retinal pS396-tau**.**

**eFigure 5.** Mapping of different retinal tau forms.

**eReference**

### Extended Methods

**Postmortem eyes from human donors**. Human eye and brain tissues collected from donor patients with premortem clinical diagnoses of MCI and AD dementia (and confirmed postmortem AD neuropathology), and age- and sex-matched CN controls (total n=75 subjects) were primarily obtained from the Alzheimer's Disease Research Center (ADRC) Neuropathology Core in the Department of Pathology (IRB protocol HS-042071) of Keck School of Medicine at the University of Southern California (USC, Los Angeles, CA). Additional eyes were obtained from the National Disease Research Interchange (NDRI, Philadelphia, PA) under approved Cedars-Sinai Medical Center IRB protocol Pro00019393. USC-ADRC, NDRI, and UCI-ADRC maintain human tissue collection protocols that are approved by their managerial committees and subject to oversight by the National Institutes of Health. Histological studies at Cedars-Sinai Medical Center were performed under IRB protocols Pro00053412 and Pro00019393. Demographic, clinical, and neuropathological information on human donors is detailed in **Table 1 and Supplementary Table 1**. Patients' identity was protected by de-identifying all tissue samples in a manner not allowing to be traced back to tissue donors.

**Clinical and neuropathological assessments**. ADRC provided the clinical and neuropathological reports on the patients' neurological examinations, neuropsychological and cognitive tests, family history, and medication lists as collected in the ADRC system using the Unified Data Set (UDS) [1]. The NDRI provided the medical history of additional patients. Most cognitive evaluations had been performed annually and, in most cases, less than one year prior to death. Cognitive testing scores from evaluations made closest to the patient's death were used for this analysis. Two global indicators of cognitive status were used for clinical assessment: the Clinical Dementia Rating (CDR scores: 0 = normal; 0.5 = very mild impairment; 1 = mild dementia; 2 = moderate dementia; or 3 = severe dementia) [2] and the Mini-Mental State Examination (MMSE scores: normal cognition = 24–30; MCI = 20–23; moderate dementia = 10–19; or severe dementia ≤ 9) [3]. In this study, the composition of the clinical diagnostic group (AD, MCI, or CN) was determined by source clinicians based on findings of a comprehensive battery of tests including neurological examinations, neuropsychological evaluations, and the aforementioned cognitive tests. To obtain a final diagnosis based on the neuropathological reports, we used the modified Consortium to Establish a Registry for Alzheimer's Disease (CERAD) [4, 5], as outlined in the National Institute on Aging (NIA)/Regan protocols with revision by the NIA and Alzheimer's Association [6]. The Aβ burden (measured as diffuse, immature, or mature plaques), amyloid angiopathy, neuritic plaques, neurofibrillary tangles (NFTs), neuropil threads (NTs), granulovacuolar degeneration, Lewy bodies, Hirano bodies, Pick bodies, balloon cells, neuronal loss, microvascular changes, and gliosis pathology were assessed in multiple brain areas, including the hippocampus (particularly the Cornu ammonis CA1, at the level of the thalamic lateral geniculate body), entorhinal cortex, superior frontal gyrus of the frontal lobe, superior temporal gyrus of the temporal lobe, superior parietal lobule of the parietal lobe, primary visual cortex (Brodmann Area-17), and visual association (Area-18) of the occipital lobe. In all cases uniform brain sampling was done by a neuropathologist.

Cerebral amyloid plaques, NFTs, and NTs were evaluated using anti–β-amyloid mAb clone 4G8 immunostaining, Thioflavin-S (ThioS) histochemical stain, and Gallyas silver stain in formalin-fixed, paraffin-embedded tissue sections. Neuropathologists (Chief, C.A.M. and Dr. Debra Hawes) provided severity scores based on semi-quantitative observations. The scale for Aβ/neuritic plaques was determined by 4G8- and/or Thioflavin-S-positive and/or Gallyas silver-positive plaques measured per 1 $mm^2$ brain area (0 = none; 1 = sparse [≤ 5 plaques]; 3 = moderate [6–20 plaques]; 5 = abundant/frequent [21–30 plaques or greater]; or N/A = not applicable), as previously described [7]; NACC NP Guidebook, Version 10, January 2014: https://naccdata.org/data-collection/forms-documentation/np-10]. Brain NFT or NT severity scoring system was derived from observed burden of these AD neuropathologic changes detected by Gallyas silver and/or Thioflavin-S staining [7-9] and measured per 1 $mm^2$ brain area. The assigned NFT or NT scores are as following: 0 = none; 1 = sparse (mild burden); 3 = moderate (intermediate burden); or 5 = frequent (severe burden). In both histochemical and immunohistochemical staining, each anatomic area of interest is assessed for the relevant pathology using the 20X objective (200X high power magnification) and representative fields are graded using a semiquantitative scale as detailed above. Validation of AD neuropathic change (ADNP), especially NTs, is performed using the 40X objectives (400X high power magnification); an average of 2 readings was assigned to each individual patient.

A final diagnosis included AD neuropathological change using an "ABC" score derived from 3 separate 4-point scales. We used the modified Aβ plaque Thal score (A0 = no Aβ or amyloid plaques; A1 = Thal phase 1 or 2; A2 = Thal phase 3; or A3 = Thal phase 4 or 5) [10]. For the NFT stage, the modified Braak staging for silver-based histochemistry or p-tau IHC was used (B0 = no NFTs; B1 = Braak stage I or II; B2 = Braak stage III or IV; or B3 = Braak stage V or VI) [11]. For the neuritic plaques, we used the modified CERAD score (C0 = no neuritic plaques; C1 = CERAD score sparse; C2 = CERAD score moderate; or C3 = CERAD score frequent) [4]. Neuronal loss, gliosis, granulovacuolar degeneration, Hirano bodies, Lewy bodies, Pick bodies, and balloon cells were all evaluated (0 = absent or 1 = present) in multiple brain areas by staining tissues with hematoxylin and eosin (H&E). Brain atrophy was evaluated (0 = none; 1 = mild; 3 = moderate; 5 = severe; or 9 = not applicable).

**Processing of eye and brain tissues**. Donor eyes were collected within an average of 8 hours after time of death and were 1) preserved in Optisol-GS media (Bausch & Lomb, 50006-OPT) and stored at 4°C for less than 24 hours; 2) fresh frozen (snap frozen; stored at -80°C); or 3) punctured once and fixed in 10% neutral buffered formalin (NBF) or 4% paraformaldehyde (PFA) and stored at 4°C. Regardless of the source of the human donor eye (USC-ADRC, UCI-ADRC or NDRI), the same tissue collection and processing methods were applied.

**Preparation of retinal strips**. Eyes that were fixed in 10% NBF or 4% PFA were dissected to create eyecups. The complete neurosensory retinas were isolated, detached from the choroid and sclera, flat mounts were prepared, and vitreous humor thoroughly removed manually, as previously described [12]. Alternatively, fresh-frozen eyes and eyes preserved in Optisol-GS were dissected, with the anterior chambers removed to create eyecups. Vitreous humor liquid was allowed to flow out and the complete neurosensory retina was isolated. Next, the vitreous gel was further thoroughly removed. Fresh-frozen retina was isolated in cold PBS with 1× Protease Inhibitor cocktail set I (Calbiochem 539131). For all flat mount retinas, the 4 topographical quadrants were defined by identifying the macula, optic disc (OD), and blood vessels [13]. Flat mount strips (~2mm wide) were prepared from superior-temporal retina, spanning diagonally from the OD to the ora serrata. Fixed retinal strips were processed for cross-sectioning. Fresh retinal strips (~5mm wide) were prepared and stored at -80°C for protein analysis. In a subset of freshly isolated donor eyes, additional ~2mm-wide strips were dissected and fixed in 4% PFA for processing of retinal cross-sections. Each strip measured approximately 2-2.5cm from the optic disc to the ora serrata. This sample preparation technique allowed for extensive and consistent access to retinal quadrants, layers, and pathological subregions.

**Paraffin-embedded retinal cross-sections**. Flat mount-derived strips were initially paraffinized using the standard techniques. Next, strips were embedded in paraffin after flip-rotating 90° horizontally. The retinal strips were sectioned (7-10μm thick) and placed on microscope slides treated with 3-aminopropyltriethoxysilane (APES, Sigma A3648). Before immunohistochemistry, the sections were deparaffinized with 100% xylene twice (10 min each), rehydrated with decreasing concentrations of ethanol (100% to 70%), and washed with distilled water followed by PBS.

**Immunofluorescent staining**. Following deparaffinization, retinal sections were incubated in blocking buffer (Dako #X0909), followed by primary antibody incubation (information provided in **Supplementary table 1**) overnight in 4°C. Retinal sections were then washed 3 times by PBS and incubated with secondary antibodies against each species (1:200, information provided in **Supplementary table 1**) for 1 hr at RT. After rinsing with PBS for 3 times, sections were mounted with Prolong Gold antifade reagent with DAPI (Thermo Fisher #P36935).

**Peroxidase-based immunostaining.** Fixed brain sections and retinal cross-sections after deparaffinization were treated with target retrieval solution (pH 6.1; S1699, DAKO) at 98 °C for 1 h and washed with PBS. Peroxidase-based immunostaining was performed. For antibodies' list and dilutions, see **Supplementary Table 1**. Prior to peroxidase-based immunostaining, the tissues were treated with 3% H2O2 for 10 min, and two staining protocols were used: (1) Vectastain Elite ABC HRP kit (Vector, PK-6102, Peroxidase Mouse IgG) according to manufacturer's instructions or (2) All Dako reagents protocol. Following the treatment with formic acid, the tissues were washed with wash buffer (Dako S3006) for 1 h, then treated with H2O2 and rinsed with wash buffer. Primary antibody (Ab) was diluted with background reducing components (Dako S3022) and incubated with the tissues overnight at 4 °C. Tissues were rinsed twice with wash buffer on a shaker and incubated for 30 min at 37 °C with secondary Ab (goat anti mouse ab HRP conjugated, DAKO Envision K4000), then were rinsed again with wash buffer. For both protocols, diaminobenzidine (DAB) substrate was used (DAKO K3468). Counterstaining with hematoxylin was performed followed by mounting with Faramount aqueous mounting medium (Dako, S3025). Routine controls were processed using identical protocols while omitting the primary antibodies to assess nonspecific labeling.

**Bielschowsky's silver staining.** Bielschowsky's silver staining. Fixed brain sections and retinal cross-sections were deparaffinized then processed for silver stain following Hito Bielshowsky OptimStainTM Kit (Hito, #HTNKS1126). Protocol was optimized for our samples. Sections were first incubated with solution-1 for 22.5 minutes at 4°C in dark humidity chamber. Meanwhile Developer solution was prepared by mixing 1mL of solution-1 with 600ul solution-2 (add 10µl solution-2 for 17-20 times until solution is clear). Portion of the final solution was set aside (1ml) and label as Developer solution. After incubation each section was rinsed for 3 times with distilled water, followed by addition of prepared mixture onto sections and incubation in dark humidity chamber at 4°C for 22.5 minutes. Slides were then quickly rinsed in distilled water. Double distilled water (50ml) and Solution 2 (200µl) were added to a coplin jar and mixed well. Slides were placed in coplin jar once incubation was completed until developing step. Solution 3 (60µl) and Developer solution (1ml) were added to an Eppendorf tube. New solution was immediately dropped on sections till fully covered. Sections were left covered in humidity chamber routinely checking on degree of color change every 30 seconds to 1 minute. After 4 minutes sections were moved to light microscope and checked for staining level. Sections should turn brown taking approximately 7.5/8minutes to achieve full staining (times may vary). Once tissue was golden brown sections were placed in coplin jar for 1 minute. Solution 4 to a 12-ml staining jar (provided by kit and slides were added to the staining jar for 3 minutes at room temperature. Slides were then rinsed in double distilled water 2 times for 2 minutes each. The slides were then dehydrated in 50% EtOH, 75% EtOH, 95% EtOH, 100% EtOH with 2 changes in each step and 3-5 minutes during each change. The slides were clear in xylene 2 times, 4 minutes each. Resinous based mounting media and coverslips were applied to each slide then allowed to dry for brightfield microscopy.

**Microscopy.** Fluorescence and bright field images were acquired using a Carl Zeiss Axio Imager Z1 fluorescence microscope with ZEN 2.6 blue edition software (Carl Zeiss MicroImaging, Inc.) equipped with ApoTome, AxioCam MRm, and AxioCam HRc cameras. Multi-channel image acquisition was used to create images with multiple channels. Images were repeatedly captured at the same focal planes with the same exposure time. Images were captured at 20×, 40×, and 63× objectives for different purposes.

**Stereological Quantification.** the fluorescence of specific signals was captured using the same setting and exposure time for each image by the Axio Imager Z1 microscope (with motorized Z-drive) with an AxioCam MRm monochrome camera (version 3.0; at a resolution of 1388 × 1040 pixels, 6.45 µm × 6.45 µm pixel size, and a dynamic range of >1:2200, which delivers low-noise images due to a Peltier-cooled sensor). Images were captured at 20x or 40x objectives, at a respective resolution of 0.25 µm. About twenty images were obtained from each retina. Acquired images were converted to gray scale and standardized to baseline using a histogram-based threshold in the Fiji ImageJ (NIH) software program (version 1.53c). For each biomarker, the total area of immunoreactivity was determined using the same threshold percentage from the baseline in ImageJ (with the same percentage threshold setting for all diagnostic groups). The images were then subjected to particle analysis to determine the immunoreactive (IR) area and or area fraction (%).

**GeoMx Digital Spatial Profiling** of total tau and phosphorylated tau (p-tau). Formalin fixed paraffin embedded human brain (A-9; frontal lobe) and retinal cross-sections from ST and IT regions (spanning from the optic disc to the ora serrata) deparaffinized before the IHC procedure, with 100% xylene twice (10 min each), rehydrated with decreasing concentrations of ethanol (100% to 70%), and washed with distilled water followed by PBS. Before fluorescence-based immunostaining were performed deparaffinized brain and retinal cross-sections were treated with target retrieval solution (pH 6.1; S1699, DAKO) at 99 °C for 1 h, washed with PBS, and then treated in formic acid 70% (ACROS) for 20 min at room temperature (RT) and washed with PBS. Subsequently, sections were treated with blocking solution (DAKO X0909) supplemented with 0.2% Triton X-100 (Sigma, T8787) prior to overnight incubation with primary Abs (for morphology markers) at 4 °C. Secondary Abs (Cy2 and Cy3) were added the following day and incubated for 1.5 hr at RT.

Selecting region of interest (ROI) by using IHC morphological markers detect proteins. For ROI selection, whole slides were stained by using three morphology markers: 1. Tissue marker –Aβ antibody (MOAB-2; NBP2-13075;

Novus; dilution 1:500) recognizes unaggregated, oligomeric and fibrillar forms of Aβ42 and unaggregated Aβ40 and does not detect APP. 2. Immune cell marker – Iba1/AIF1 antibody recognizes microglial/macrophage (20A12.1; 970896; EMD Milipore; dilution 1:300). 3. DNA marker – Syto 13. For all subjects' groups (AD, MCI and NC) the same ROI dimensions were selected from the retinal C, M and f subregions, and in total three ROI per retinal cross section. Also, the same ROI dimensions were selected from each brain section, and in total two ROI per brain section.

For the tau module, a panel of digital spatial profiler (DSP) barcoded antibodies were used in GeoMx Protein assays. The DSP barcode was conjugated to the antibody with photocleavable linker. The antibodies were against total tau and p-tau (S214; T231; S199; S396 and S404)

Following staining the slides were imaged and profiled using GeoMx® DSP, and tissues were exposed to UV light in the selected ROI. The UV cleaved the DSP barcode from the antibody. The protein expression level was collected and quantifying directly with digital counting of released barcodes using the nCounter® Analysis System. Raw data were analyzed with DSP Data Analysis Suite (DSPDA), then the data were normalized to housekeeping protein GAPDH that was detected with the Tau module in the same experiment.

**Statistical Analysis.** GraphPad Prism version 8.3.0 (GraphPad Software) was used for the analyses. Three or more group comparisons were analyzed using one-way ANOVA followed by Tukey's multiple comparison test. Two-group comparisons were analyzed using a two-tailed unpaired Student t-test. The statistical association between two or more variables was determined using Pearson's correlation coefficient (r) test (Gaussian-distributed variables; GraphPad Prism). Pearson's r indicates the direction and strength of the linear relationship between two variables. Required sample sizes for two group (differential mean) comparisons were calculated using the nQUERY t-test model, assuming a two-sided $\alpha$ level of 0.05, 80% power, and unequal variances, with the means and common standard deviations for the different parameters. Results are expressed as means ± SDs. A P value less than 0.05 is considered significant.

**Supplementary Table 1. List of human donors in this study.**

| Diagnosis | Sex | Race | Age at death | Thal-A | Braak-B | CERAD-C | CAA Score | Co-morbid. [LB/ASVD] | Braak Stage | CDR Score | MMSE Score | APOE status | Study type |
|---|---|---|---|---|---|---|---|---|---|---|---|---|---|
| AD1 | F | W | 93 | 2 | 3 | 3 | 1 | −/+ | V | 3 | n.a. | n.a. | IHC |
| AD2 | M | H | 97 | 3 | 2 | 3 | 1 | −/+ | III | n.a. | n.a. | n.a. | IHC |
| AD3 | M | W | 79 | 3 | 3 | 3 | 1.5 | −/+ | V | n.a. | n.a. | n.a. | IHC |
| AD4 | F | W | 87 | 3 | 3 | 3 | n.a. | −/+ | V-VI | n.a. | n.a. | n.a. | IHC |
| AD5 | M | W | 88 | 2 | 3 | 2 | 1 | −/+ | V-VI | 1 | 18 | e3/e4 | IHC |
| AD6 | M | W | 77 | 3 | 3 | 3 | 1 | −/− | VI | 2 | n.a. | e3/e4 | IHC |
| AD7 | F | W | 66 | 3 | 3 | 3 | 0 | −/+ | VI | 3 | n.a. | e3/e3 | IHC |
| AD8 | F | H | 99 | 3 | 2 | 3 | 1.5 | −/− | IV | n.a. | n.a. | n.a. | IHC |
| AD9 | F | H | 81 | 3 | 3 | 3 | 1.5 | −/+ | V-VI | 3 | n.a. | e3/e3 | IHC/GeoMx |
| AD10 | F | W | 90 | 3 | 3 | 3 | 1 | −/+ | V-VI | 3 | n.a. | e3/e4 | IHC/GeoMx |
| AD11 | M | W | 90 | 3 | 2 | 3 | 1 | −/+ | III-IV | 3 | 1 | e3/e4 | IHC/GeoMx |
| AD12 | M | W | 90 | 3 | 3 | 3 | 1 | −/+ | VI | 3 | n.a. | n.a. | IHC/GeoMx |
| AD13 | F | W | 90 | 1 | 3 | 3 | 1 | −/+ | V | 2 | 9 | n.a. | IHC/GeoMx |
| AD14 | M | W | 79 | 2 | 2 | 2 | 2 | −/+ | V | 0.5 | 24 | n.a. | IHC |
| AD15 | F | H | 92 | 2 | 2 | 2 | 2 | −/+ | III | 3 | n.a. | e3/e3 | IHC |
| AD16 | M | W | 88 | 3 | 3 | 3 | 1.5 | −/+ | V-VI | 1 | 16 | e2/e3 | IHC/GeoMx |
| AD17 | F | B | 94 | 3 | 3 | 3 | 0 | −/+ | V-VI | 3 | n.a. | e3/e3 | IHC |
| AD18 | F | n.a. | 87 | n.a. | n.a. | n.a. | n.a. | n.a./n.a. | n.a. | n.a. | n.a. | n.a. | IHC |
| AD19 | F | n.a. | 70 | n.a. | n.a. | n.a. | n.a. | n.a./n.a. | n.a. | n.a. | n.a. | n.a. | IHC |
| AD20 | M | A | 81 | 3 | 3 | 3 | 1 | −/− | V | 1 | n.a. | e4/e4 | IHC |
| AD21 | M | W | 79 | 3 | 1 | 3 | n.a. | +/+ | I | n.a. | n.a. | n.a. | IHC |
| AD22 | M | W | 92 | 2 | 3 | 3 | 0 | −/+ | VI | 3 | 2 | e3/e3 | IHC |
| AD23 | M | W | 66 | 3 | 3 | 3 | 1.5 | −/n.a. | V | 3 | n.a. | n.a. | IHC/GeoMx |
| AD24 | M | A | 40 | 3 | 2 | 3 | 1 | −/− | III-IV | 3 | n.a. | e3/e3 | IHC |
| AD25 | F | A | 88 | 2 | 3 | 3 | 1.5 | −/+ | V | 3 | n.a. | n.a. | IHC |
| AD26 | F | W | 100 | 2 | 3 | 3 | 1 | −/+ | V-VI | 2 | 16 | n.a. | IHC |

| | | | | | | | | | | | | | |
|---|---|---|---|---|---|---|---|---|---|---|---|---|---|
| AD27 | F | W | 86 | 3 | 3 | 2 | 1 | −/+ | V-VI | 2 | n.a. | e3/e4 | IHC/ Geo Mx |
| AD28 | F | W | 85 | 3 | 3 | 3 | 1.5 | −/+ | V-VI | 3 | n.a. | e3/e3 | IHC/ Geo Mx |
| AD29 | F | B | 93 | 2 | 2 | 2 | 1.5 | +/+ | III-IV | 3 | n.a. | n.a. | IHC |
| AD30 | F | W | 76 | 3 | 3 | 3 | 2 | −/+ | V | 1 | 26 | e3/e4 | IHC |
| AD31 | F | A | 93 | 3 | 2 | 2 | 0 | −/+ | III-IV | 3 | n.a. | e3/e3 | IHC |
| AD32 | F | W | 97 | n.a. | n.a. | n.a. | 0 | −/+ | IV | n.a. | 4 | n.a. | IHC |
| AD33 | F | W | 63 | n.a. | n.a. | n.a. | 0 | −/+ | V | n.a. | 9 | n.a. | IHC |
| AD34 | F | W | 65 | n.a. | n.a. | n.a. | 2 | −/+ | V-VI | n.a. | n.a. | n.a. | IHC |
| MCI1 | M | W | 97 | 2 | 3 | 3 | 1 | −/+ | V | 0.5 | n.a. | e3/e3 | IHC |
| MCI2 | M | H | 80 | 3 | 3 | 2 | n.a. | −/+ | V-VI | 3 | n.a. | e3/e3 | IHC/ Geo Mx |
| MCI3 | F | B | 94 | 3 | 1 | 1 | 0 | n.a./− | I-II | 0.5 | 23 | e3/e3 | IHC/ Geo Mx |
| MCI4 | F | W | 89 | 1 | 2 | 2 | 1 | −/+ | III-IV | 0.5 | 21 | e3/e3 | IHC/ Geo Mx |
| MCI5 | F | W | 93 | 3 | 2 | 2 | 2 | −/+ | IV | 3 | n.a. | e3/e3 | IHC |
| MCI6 | M | W | 93 | 2 | 2 | 0 | 0 | −/+ | 0 | 3 | n.a. | e2/e3 | IHC/ Geo Mx |
| MCI7 | F | W | 86 | 3 | 1 | 3 | 0 | −/+ | I-II | 0 | 26 | e3/e4 | IHC |
| MCI8 | M | W | 88 | 1 | 2 | 2 | 0 | −/+ | III | 0 | 24 | n.a. | IHC/ Geo Mx |
| MCI9 | F | W | 80 | n.a. | n.a. | n.a. | 0 | +/n.a. | IV | n.a. | n.a. | n.a. | IHC |
| MCI10 | F | W | 87 | 3 | 3 | 3 | 1.5 | −/+ | V-VI | 3 | n.a. | e3/e3 | IHC/ Geo Mx |
| MCI11 | F | W | 98 | n.a. | n.a. | n.a. | 1.5 | −/+ | V | 2 | 15 | n.a. | IHC |
| NC1 | F | W | 93 | n.a. | n.a. | n.a. | n.a. | n.a. | n.a. | n.a. | n.a. | n.a. | IHC |
| NC2 | F | W | 86 | n.a. | n.a. | n.a. | n.a. | n.a. | n.a. | n.a. | n.a. | n.a. | IHC |
| NC3 | M | W | 78 | n.a. | n.a. | n.a. | n.a. | n.a. | n.a. | n.a. | n.a. | n.a. | IHC |
| NC4 | M | H | 76 | n.a. | n.a. | n.a. | n.a. | n.a. | n.a. | n.a. | n.a. | n.a. | IHC |
| NC5 | M | W | 95 | 1 | 1 | 1 | 0 | −/− | I | 0 | n.a. | e3/e3 | IHC/ Geo Mx |
| NC6 | F | W | 88 | n.a. | n.a. | n.a. | n.a. | n.a. | n.a. | n.a. | n.a. | n.a. | IHC |
| NC7 | M | H | 81 | 3 | 2 | 2 | 0 | −/+ | I-II | 0 | 23 | e3/e4 | IHC/ Geo Mx |
| NC8 | F | W | 99 | 1 | 2 | 1 | 0 | −/− | III | 0 | n.a. | e3/e3 | IHC/ Geo Mx |

| | | | | | | | | | | | | | |
|---|---|---|---|---|---|---|---|---|---|---|---|---|---|
| NC9 | F | W | 92 | n.a. | n.a. | n.a. | 0 | −/+ | I-II | n.a. | 25 | n.a. | IHC/ Geo Mx |
| NC10 | M | W | 69 | n.a. | n.a. | n.a. | 0 | −/+ | 0 | n.a. | 28 | n.a. | IHC/ Geo Mx |
| NC11 | F | W | 91 | n.a. | n.a. | n.a. | 0 | −/+ | III | n.a. | 29 | n.a. | IHC |
| NC12 | M | W | 77 | n.a. | n.a. | n.a. | n.a. | n.a. | n.a. | n.a. | n.a. | n.a. | IHC |
| NC13 | M | W | 73 | n.a. | n.a. | n.a. | n.a. | n.a. | n.a. | n.a. | n.a. | n.a. | IHC |
| NC14 | M | W | 84 | n.a. | n.a. | n.a. | n.a. | n.a. | n.a. | n.a. | n.a. | n.a. | IHC |
| NC15 | M | W | 70 | n.a. | n.a. | n.a. | n.a. | n.a. | n.a. | n.a. | n.a. | n.a. | IHC |
| NC16 | F | W | 95 | 3 | 3 | 2 | 1 | −/+ | V | 0 | 30 | e3/e3 | IHC/ Geo Mx |
| NC17 | F | W | 93 | 3 | 2 | 3 | 1 | −/+ | III-IV | 1 | n.a. | e2/e3 | IHC/ Geo Mx |
| NC18 | F | H | 85 | 2 | 1 | 2 | 0 | −/+ | I-II | 0 | 30 | e3/e3 | IHC/ Geo Mx |
| NC19 | M | B | 80 | n.a. | n.a. | n.a. | n.a. | n.a. | n.a. | n.a. | n.a. | n.a. | IHC |
| NC20 | M | W | 58 | n.a. | n.a. | n.a. | n.a. | n.a. | n.a. | n.a. | n.a. | n.a. | IHC |
| NC21 | M | W | 75 | n.a. | n.a. | n.a. | n.a. | n.a. | n.a. | n.a. | n.a. | n.a. | IHC |
| NC22 | M | W | 87 | n.a. | n.a. | n.a. | n.a. | n.a. | n.a. | n.a. | n.a. | n.a. | IHC |
| NC23 | M | W | 75 | n.a. | n.a. | n.a. | n.a. | n.a. | n.a. | n.a. | n.a. | n.a. | IHC |
| NC24 | F | W | 76 | n.a. | n.a. | n.a. | n.a. | n.a. | n.a. | n.a. | n.a. | n.a. | IHC |
| NC25 | F | W | 75 | n.a. | n.a. | n.a. | n.a. | n.a. | n.a. | n.a. | n.a. | n.a. | IHC |
| NC26 | F | W | 76 | n.a. | n.a. | n.a. | n.a. | n.a. | n.a. | n.a. | n.a. | n.a. | IHC |
| NC27 | F | W | 77 | n.a. | n.a. | n.a. | n.a. | n.a. | n.a. | n.a. | n.a. | n.a. | IHC |
| NC28 | F | W | 71 | n.a. | n.a. | n.a. | n.a. | n.a. | n.a. | n.a. | n.a. | n.a. | IHC |
| NC29 | F | B | 73 | n.a. | n.a. | n.a. | n.a. | n.a. | n.a. | n.a. | n.a. | n.a. | IHC |
| NC30 | F | W | 95 | n.a. | n.a. | n.a. | n.a. | −/− | I | n.a. | n.a. | n.a. | IHC/ Geo Mx |

AD, Alzheimer's disease dementia; MCI, mild cognitive impairment; NC, normal cognition; F, female; M, male; A, Asian; B, Black; H, Hispanic; W, White; IHC, Immunohistochemistry; A, Aβ plaque score modified from Thal; B, NFT stage modified from Braak; C, Neuritic plaque score modified from CERAD; CAA, Cerebral amyloid angiopathy; LB, Lewy bodies; ASVD. Atherosclerosis; CDR, Clinical dementia rating; MMSE, Mini-Mental State Examination; n.a., not available; +: present; –: none; APOE, apolipoprotein alleles.

**Supplementary Table 2.** List of antibodies.

| Antibodies or Reagents | Source Species | Dilution | Application | Commercial Source | Catalog. # |
|---|---|---|---|---|---|
| *Primary antibody* | | | | | |
| MC1 mAb | Mouse | 1:200 | IF, DAB | - | - |
| Anti-tau T22 pAb | Rabbit | 1:200 | IF | - | - |
| Tau Monoclonal Antibody (HT7) | Mouse | 1:500 | IF | ThermoFisher | MN1000 |
| Purified anti-Tau, 1-100 Antibody (43D) | Mouse | 1:40 | IF | Biolegend | 816601 |
| Phospho-tau (Thr212, Ser214) mAb (AT100) | Mouse | 1:100 | IF, DAB | ThermoFisher | MN1060 |
| Phospho-tau (Ser202, Thr205) mAb (AT8) | Mouse | 1:250 | IF, DAB | ThermoFisher | MN1020 |
| Phospho-tau (Ser396) pAb | Rabbit | 1:1500 | IF, DAB | Anaspec | AS-54977 |
| Tau phos Ser396/Ser404 mAb (PHF-1) | Mouse | 1:200 | IF | - | - |
| Cit$_{209}$tau | Mouse | 1:5000 | IF | - | - |
| βIII-tubulin mAb | Mouse | 1:1000 | IF | Abcam | Ab78078 |
| PAD4 | Mouse | 1:100 | IF | Abcam | Ab128086 |
| Iba1 pAb | Rabbit | 1:200 | IF | FUJIFILM | 019-19741 |
| *Secondary antibody* | | | | | |
| Cy2 (anti-Rabbit) | Donkey | 1:200 | IF | Jackson ImmunoResearch Laboratories | |
| Cy3 (anti-rabbit, anti-goat, anti-mouse) | Donkey | 1:200 | IF | Jackson ImmunoResearch Laboratories | |
| Cy5 (anti-mouse) | Donkey | 1:200 | IF | Jackson ImmunoResearch Laboratories | |

Abbreviation: IF – immunofluorescence; DAB - peroxidase-based immunohistochemistry visualized with DAB substrate; pAb – polyclonal antibody; mAb – monoclonal antibody.

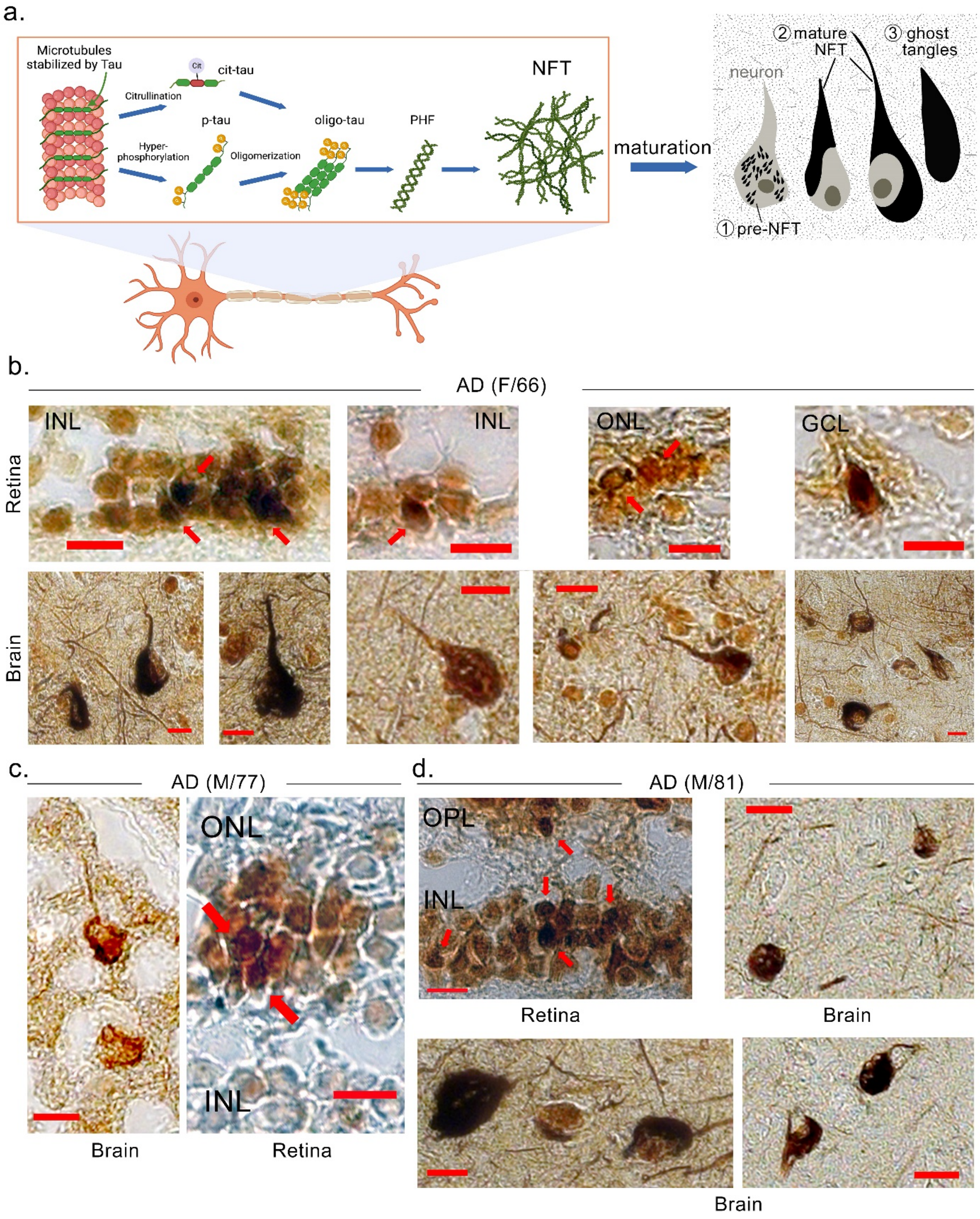


**Supplementary Figure 1.** Bielschowsky silver staining for brain and retinal NFTs. a. Illustration of different forms of abnormal tau created by using Biorender.com. b-d. Bielschowsky silver staining for brain and retinal NFTs from different AD patients. Scale bars=10μm.

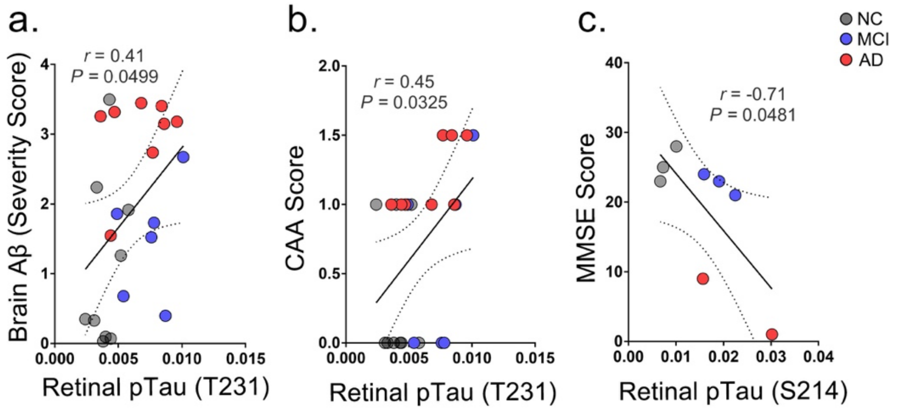


**Supplementary Figure 2.** Additional correlations between GeoMx® DSP of retinal p-tau and brain pathology. **a-c.** Pearson's coefficient (*r*) correlation between **a.** retinal p-tau at T231 vs. brain Aβ burden (n=23) **b.** retinal p-tau at T231 vs. cerebral amyloid angiopathy severity score (n=23), and **c.** retinal p-tau at S214 vs. cognitive score by Mini-Mental State Examination (MMSE, n=8).

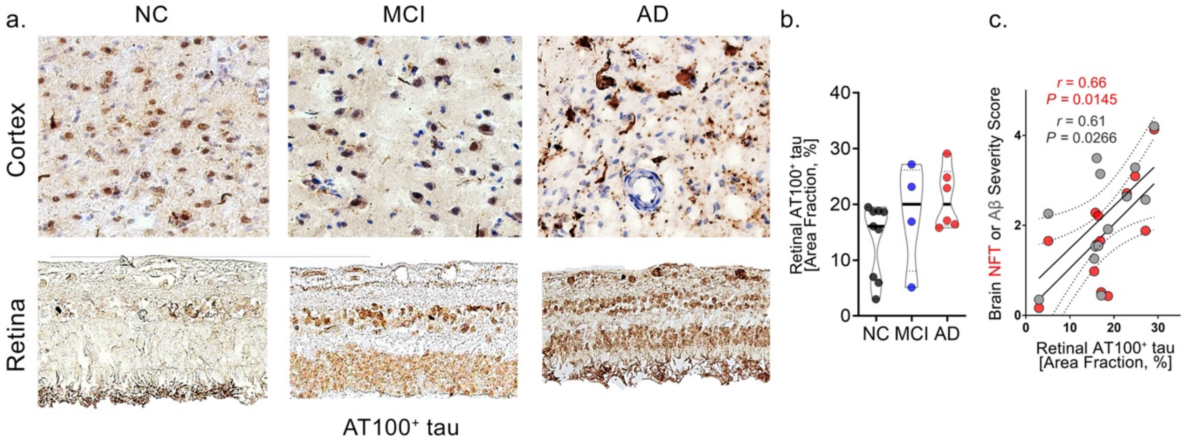


**Supplementary Figure 3.** Retinal AT100$^+$p-tau burden and association with brain pathology. **a.** Representative images of peroxidase-based staining of AT100$^+$p-tau (T202/S214) on retinal cross-sections from MCI, AD patients and normal cognition (NC) controls. **b.** Quantitative analysis of retinal AT100$^+$p-tau immunoreactivity (IR) (n=46 in total). **c.** Pearson's coefficient (*r*) correlation between retinal AT100+p-tau vs. brain NFTs burden (n=13) or Aβ burden (n=13). Data from individual donors (circles) as well as group means ± SDs are shown.

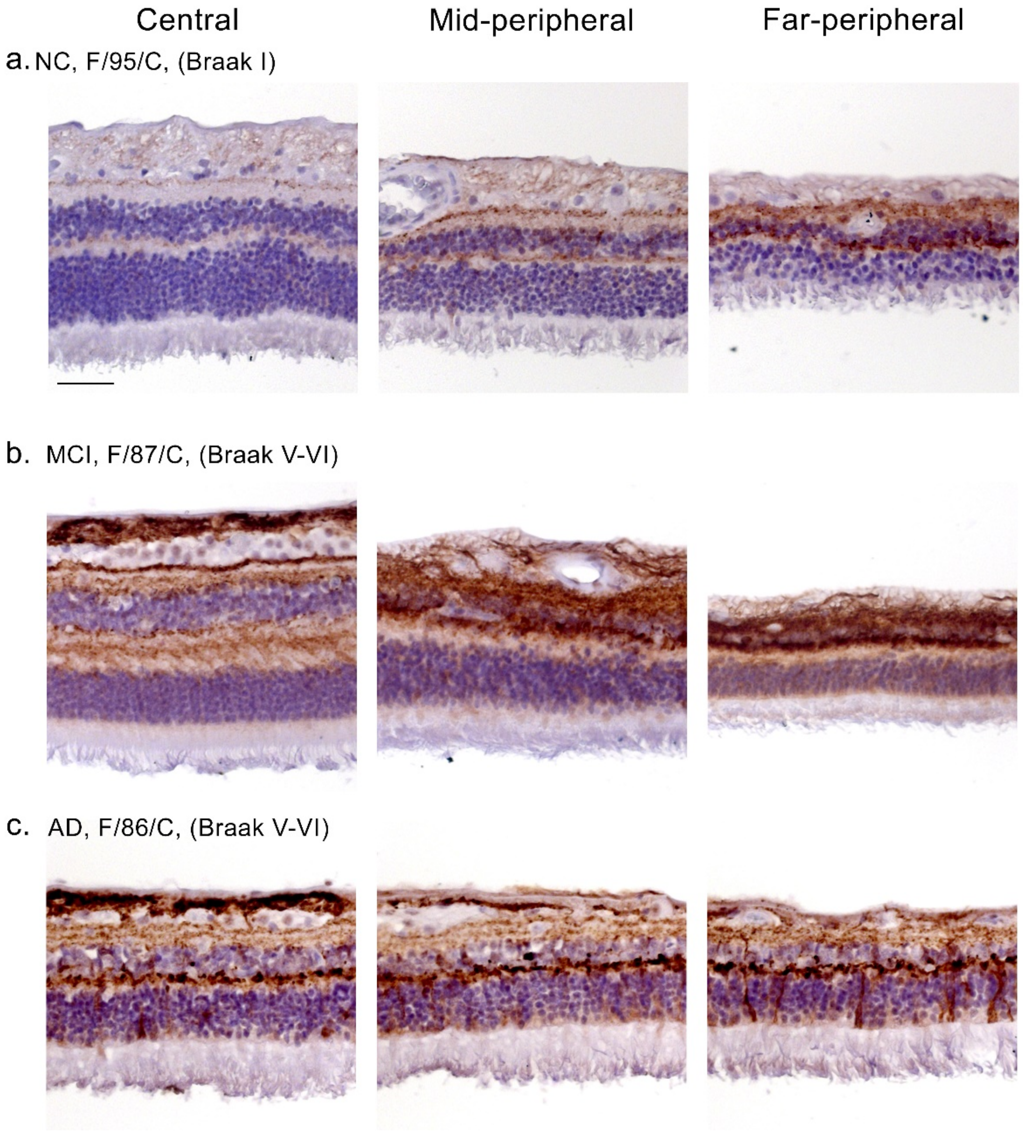


**Supplementary Figure 4.** Regional difference of retinal pS396-tau**. a-c.** Representative images of peroxidase-based staining of pS396$^{+}$p-tau on retinal cross-sections from MCI, AD patients and normal cognition (NC) controls grouped by central, mid-peripheral, and far-peripheral retinal regions.

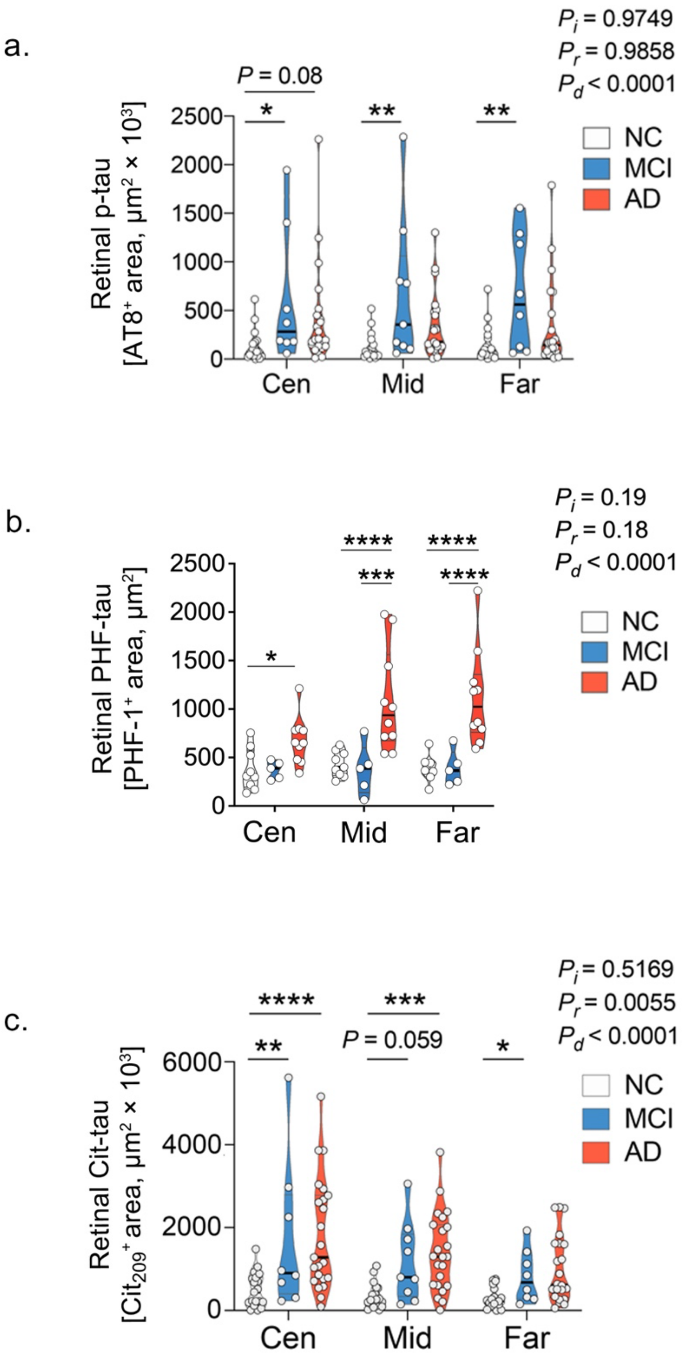


**Supplementary Figure 5.** Mapping of different retinal tau forms. **a-c**. Mapping of $AT8^+$, PHF-, and $cit_{209}$-tau forms in central, mid-, and far-peripheral retinal regions. Data from individual donors (circles) as well as group means ± SDs are shown. *$p < 0.05$, **$p < 0.01$, ***$p < 0.001$, ****$p < 0.0001$, by one-way ANOVA with Tukey's post-hoc multiple comparison test.
